# Evolutionary graph theory on rugged fitness landscapes

**DOI:** 10.1101/2023.05.04.539435

**Authors:** Yang Ping Kuo, Oana Carja

**Affiliations:** Computational Biology Department, School of Computer Science, Carnegie Mellon University, Pittsburgh, PA, USA; Joint Carnegie Mellon University-University of Pittsburgh Ph.D. Program in Computational Biology

**Keywords:** evolutionary graph theory, spatial structure, population structure, rugged fitness landscapes, stochastic tunneling

## Abstract

Spatially-resolved datasets are revolutionizing knowledge in molecular biology, yet are under-utilized for questions in evolutionary biology. To gain insight from these large-scale datasets of spatial organization, we need mathematical representations and modeling techniques that can both capture their complexity, but also allow for mathematical tractability. Specifically, it is hard to link previous deme-based or lattice-based models with datasets exhibiting complex patterns of spatial organization and the role of heterogeneous population structure in shaping evolutionary dynamics is still poorly understood. Evolutionary graph theory utilizes the mathematical representation of networks as a proxy for population structure and has started to reshape our understanding of how spatial structure can direct evolutionary dynamics. However, previous results are derived for the case of a single mutation appearing in the population. Complex traits arise from interactions among multiple genes and these interaction can result in rugged fitness landscapes, where evolutionary dynamics can vastly differ from the dynamics of stepwise fixation. Here, we develop a unifying theory of how heterogenous population structure shapes evolutionary dynamics on rugged fitness landscapes. We show that even a simple extension to a two- mutational landscape can exhibit evolutionary dynamics not observed in deme-based models and that cannot be predicted using previous single-mutation results. We also show how to link these models to spatially-resolved datasets and build the networks of the stem cell niches of the bone marrow. We show that these cellular spatial architectures reduce the probability of neoplasm initiation across biologically relevant mutation rate and fitness distributions.

## Introduction

The recent influx of spatially-resolved studies is starting to give us unprecedented access to molecular and cellular patterns of spatial organization (Xia et al., 2019; Galeano Niño et al., 2022; Ratz et al., 2022; Rodriques et al., 2019; Zhao et al., 2022; Erickson et al., 2022; Lomakin et al., 2022; Peurla et al., 2023; Gomariz et al., 2018; Rindone et al., 2021; Rendeiro et al., 2021; Goltsev et al., 2018; Mondragón-Palomino et al., 2022). While classic models in population genetic theory have been extraordinarily important for producing initial testable predictions about the role of space and structure in evolution, most previous modeling approaches represent spatial structure as a small number of distinct patches, symmetrically connected by migration corridors (Wright, 1943; Kimura and Weiss, 1964; Carja et al., 2014; Maruyama, 1970a; Slatkin, 1981; Whitlock and Barton, 1997; Whitlock, 2003; Wakeley, 2000; Durrett and Levin, 1994; Whitlock et al., 1995; Petkova et al., 2016; Gutenkunst et al., 2009; Jackson et al., 2020). This makes these models analytically tractable, but it also makes their predictions hard to link to these emerging datasets, characterized by large amounts of spatial heterogeneity. Beyond simple mathematical convenience, the level of spatial granularity incorporated in a model fundamentally shapes understanding of mechanism and the level of detail that can be ignored.

In recent years, evolutionary graph theory has started to reshape our understanding of how spatial structure can direct evolutionary dynamics (Lieberman et al., 2005; Ohtsuki and Nowak, 2006; Santos et al., 2006; Ohtsuki et al., 2006, 2007; Poncela et al., 2007; Allen et al., 2013; Maciejewski et al., 2014; Jiang et al., 2014; Leventhal et al., 2015; Kuo et al., 2021; Kuo and Carja, 2021; Su et al., 2022). Using tools from network theory, these models allow the tuning of the heterogeneity of spatial structure, beyond what is possible with deme-based or lattice-based models. The nodes of the graph represent individuals in the population and edges are proxies for the local pattern of replacement and substitution. Nodes can also be interpreted as genetically homogeneous subpopulations where genetic drift is ignored and a new beneficial mutation arriving in this node subpopulation is assumed to fix immediately. By incorporating heterogeneity in spatial structure, these models have been shown to greatly extend the range of possible evolutionary outcome of a population, beyond what was previously observed in lattice-based or symmetric deme-based models and either boost the selective benefit of new mutations appearing in a population or, reversely, suppress the spread of deleterious mutants (Adlam et al., 2015; Hindersin and Traulsen, 2015; Kuo et al., 2021; Allen et al., 2021; Tkadlec et al., 2020).

The main obstacle to progress in this area has been the analytical difficulty of multidimensional stochastic models and previous results are restricted to studying times and probabilities of fixation for one single mutation appearing in the population. However, complex traits arise from interactions between multiple gene products. In some cases, the evolution of complex traits involves the repeated accumulation of several beneficial mutations, analogous to climbing a hill in the mutation landscape (Jacob, 1977). In other cases, to reach a complex trait or a better fitness peak requires the accumulation of multiple individually neutral or even deleterious mutations and having to cross a fitness valley (Wright et al., 1932). Empirical fitness landscapes, from *Saccharomyces cerevisiae* (Kvitek and Sherlock, 2011), RNA viruses (Burch and Chao, 1999), xenobiotic-degrading enzymes (Yang et al., 2019), molecular activity experiments (Sarkisyan et al., 2016; Jiménez et al., 2013) or tumor sequencing efforts (Wood et al., 2007; Vogelstein et al., 2013; Huang, 2013; Rogers et al., 2018; Acar et al., 2020; Salehi et al., 2021) provide overwhelming evidence that rugged landscapes are prevalent in evolving populations.

A large population can cross wide fitness valleys remarkably quickly, suggesting valley-crossing dynamics is common, even when mutations that directly increase fitness are also available (Weissman et al., 2009; Weinreich and Chao, 2005; Jain and Krug, 2007). Moreover, under elevated mutation rates, crossing fitness valleys occurs mostly through stochastic tunneling, not through sequential fixation (Nowak et al., 2002; Komarova et al., 2003; Iwasa et al., 2004; Weissman et al., 2009). Deme-based and lattice-based spatial models have previously shown that spatial interactions can accelerate the crossing of fitness valleys and plateaus, when the intermediate mutants are neutral and disadvantageous, while, at the same time, delaying the evolution of complex traits with advantageous intermediates (Komarova, 2014; Bitbol and Schwab, 2014; Durrett and Moseley, 2015). However, in contrast to graph spatial topologies, these spatial structures only affect valley crossing through changes to the extinction time of the intermediate mutants, and do not affect their establishment and fixation probability (Pollak, 1966; Maruyama, 1970b; Lande, 1979). The role of network topologies in shaping multi-mutational dynamics, in particular stochastic tunneling and fitness valley crossing, has yet to be explored.

Here, we develop a unified theory of fitness landscape crossing for network-structured populations, beyond single mutation dynamics. We study the role of heterogeneous population structure in shaping rates of landscape crossing and show that previous single-mutation frameworks can only be used to predict sequential fixation dynamics, in the limit of small mutation rates.

In agreement with previous lattice and deme-based models (Komarova, 2014; Bitbol and Schwab, 2014), our results show that heterogenous spatial structure can affect fitness landscape crossing by allowing intermediate mutants to persist for longer, until the final beneficial mutation occurs. However, in stark contrast with previous models, these population structures also change the probability that the deleterious and neutral intermediates go extinct and this difference leads to a wider range of observed evolutionary behavior. We compare rates of fitness landscape crossing across well-studied families of graphs and discuss intuitive guiding principles for selecting network topologies for evolutionary optimization, when the ruggedness of the optimization landscape is known. We further define enhancers of selection networks as starting population structures that optimize for the ability to accelerate crossing over all fitness landscapes. If the ruggedness of the landscape is known, networks that specialize in a specific landscape can overall perform better than enhancers of crossing and our analytic results can be easily used to design and select the ideal network topology.

We also show how to apply these types of network-based approaches to large-scale datasets of spatial organization. We discuss the cellular networks of the stem cell niches in the bone marrow and apply our evolutionary model to study rates of mutation accumulation and leukemia initiation. Our results show that these cellular spatial architectures reduce the probability of neoplasm initiation across biologically relevant mutation rate and fitness distributions, compared to well-mixed populations. However, we show that there exists a threshold mutation rate (under exposure to carcinogens, for example) above which the bone marrow structure shifts from a suppressor into an amplifier of neoplasm initiation. More broadly, since the accumulation of mutations is important not only for cancer initiation, progression and metastasis (Friend et al., 1986; Kandoth et al., 2013; Tomasetti and Vogelstein, 2015; Loeb and Loeb, 2000; Zehir et al., 2017; Bolton et al., 2020; Samstein et al., 2019), but also microbial adaptation and evolution of antibacterial resistance (Wistrand-Yuen et al., 2018; Ogbunugafor and Eppstein, 2016; Hall et al., 2019), our results provide the groundwork for further exploration of the role of heterogeneous spatial topology for shaping rates of mutation accumulation in both engineering and biological settings.

### Model

We consider an asexual population of *N* haploid individuals and study the process by which this population acquires a beneficial trait that requires mutations at multiple loci. This process is usually referred to as the ‘K-hit’ mutational process, and the simplest case, *K* = 2 is illustrated in **Figure 1**. We consider the *K* = 2 case and assume that, at time *t* = 0, a single first mutant, with reproductive fitness (1 + *s*), appears in a population consisting of only wild-type individuals of fitness equal to one. Depending on the sign of the selection coefficient *s*, this first mutation can be beneficial, neutral or deleterious. With mutation rate *µ*, individuals carrying the first mutation can acquire a second mutation assumed to be extremely beneficial, such that fixation of double-mutant individuals is guaranteed with probability one. Therefore, the probability that all the individuals acquire the beneficial trait is equal to the probability of acquiring the second mutation. This model easily generalizes to the *K−*hit process with a *K >* 2, by starting at *k* = *K* mutations, calculating the probability of acquiring a single (*k −* 1) mutant, and working backwards to *k* = 1. Similarly, to generalize to the case when the second mutation is not guaranteed to fix, we need only multiply by the fixation probability of the second mutation, which can be written using previous results (Kuo et al., 2021).

**Figure 1:**
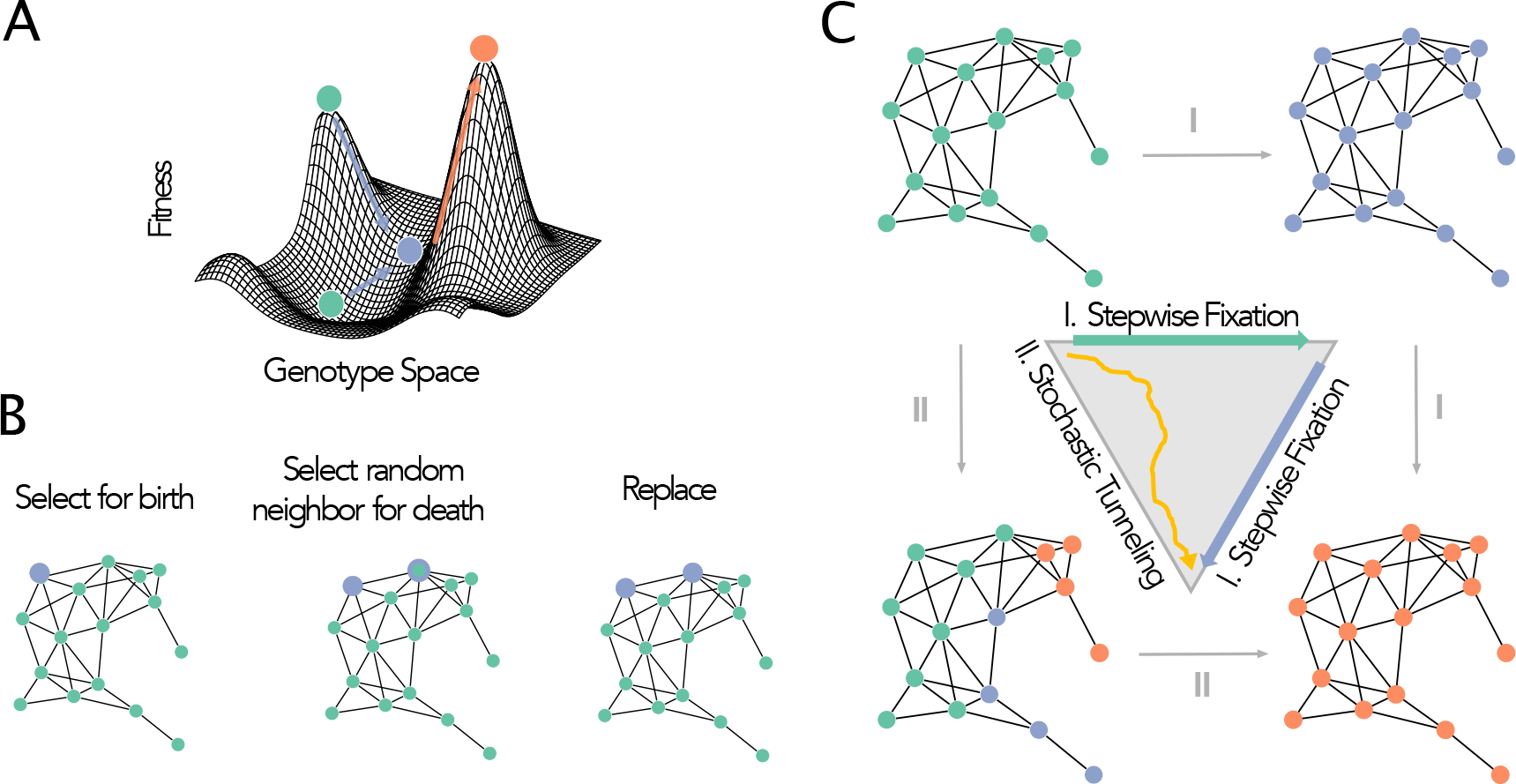
Illustration of the model. A. We represent complex mutational landscapes by using a two-step mutational process. The first mutation can be beneficial, neutral, or deleterious. The second mutation is assumed to be extremely beneficial, such that fixation is guaranteed. **B.** We illustrate the Bd (Birth-death) update rules. **C.** There are two ways for the second mutation to fix in the population. The first is through step-wise fixation, where the intermediate mutant reaches fixation before the second mutation occurs (I). The second is through stochastic tunneling, where the second mutation is acquired before fixation of the intermediate (II).

For large enough mutation rates *µ*, the second mutant will appear and spread in the population before the fixation of the first mutant and the population will transition from all wild-type to all two-mutant individuals, without stepwise fixation and without ever visiting a population fixed on the intermediate mutant (**Figure 1C**). This process is called stochastic tunneling (Iwasa et al., 2004) and it is particularly likely when the population size is large and when the intermediate mutant is deleterious, in other words, when the expected waiting time for its fixation is very long.

To represent heterogeneous population spatial structure, we use unweighted and undirected graphs, where each node in the graph represents one individual in the population and edges are proxies for the local pattern of replacement and substitution. We study the role of population structure in shaping rates of fitness landscape crossing by analyzing how graph properties determine the probability that the population reaches the fitness peak. We analyze the probability and the time it takes for the population to traverse through genotype space and arrive at the higher fitness peak as a function of the population size *N*, the mutation rate *µ*, the intermediate mutant selection coefficient *s*, and, in particular, the statistical properties of the population spatial structure.

To study graph properties independently of one another, we use graph generation algorithms and systematically tune network parameters, as described in more detail in (Kuo et al., 2021). We analyze a wide variety of graph families including preferential attachment graphs (Barabási and Albert, 1999), bipartite graphs (Asratian et al., 1998), detour graphs (Möller et al., 2019), star-like graph (Tkadlec et al., 2019), small world networks (Watts and Strogatz, 1998) used to model properties of social networks, and random geometric networks (Waxman, 1988; Penrose et al., 2003), mathematical representations of populations embedded in Euclidean space. We also introduce regular island graphs which are created by adding random edges to two separate *k*-regular graphs.

We write analytic predictions for the rate of fitness valley crossing as a function of model parameters and compare these analytic predictions with results from Monte-Carlo simulations, using at least 10^7^ simulations. These analytic results allow us to generalize our insights beyond the graph families discussed here. To simulate the evolutionary trajectories of the population, at time *t* = 0, we introduce one intermediate mutant on a random node of the network and use a Moran Birth-death (Bd) process to track mutant frequency changes (**Figure 1B**). The birth-death process assumes that at each time step, an individual from the population is selected to reproduce proportional to fitness and a random neighbor node is selected to die, leaving an unoccupied node for the offspring of the reproducing node. This is the essential difference that allows us to study the role of local population structure, in comparison to a well-mixed population: in a well-mixed population, a node from the entire network would be randomly selected for death. Each time an intermediate mutant is selected to reproduce, with probability *µ*, it can acquire the second beneficial mutation. The simulation ends when either the intermediate mutant dies out or the second mutation is acquired. The fitness landscape crossing probability is then determined by the fraction of simulations that end with acquisition of the second mutation. Landscape crossing time is determined by averaging over the time it takes for the beneficial trait to fix, given fixation.

## Results

Under a sequential fixation regime, the fitness valley crossing probability is the same as the probability of fixation of the first mutant and the role of the population structure can be understood using previous single-mutation results (**Figure 2A**). For example, with a spatial structure that amplifies selection, a beneficial first mutant has a higher rate of fitness landscape crossing compared to that of a well-mixed population, while the reverse is true for deleterious mutations. The transition between these two regimes occurs exactly at neutrality, *Ns* = 0. The population structure thus reshapes the fitness landscape into a different, effective fitness landscape. When climbing towards a fitness peak, the intermediate mutations appear more beneficial for a spatial amplifier, making the fitness peak sharper. When crossing a fitness valley, the intermediate mutations become more deleterious, making the fitness valley steeper (**Figure 2A**).

**Figure 2:**
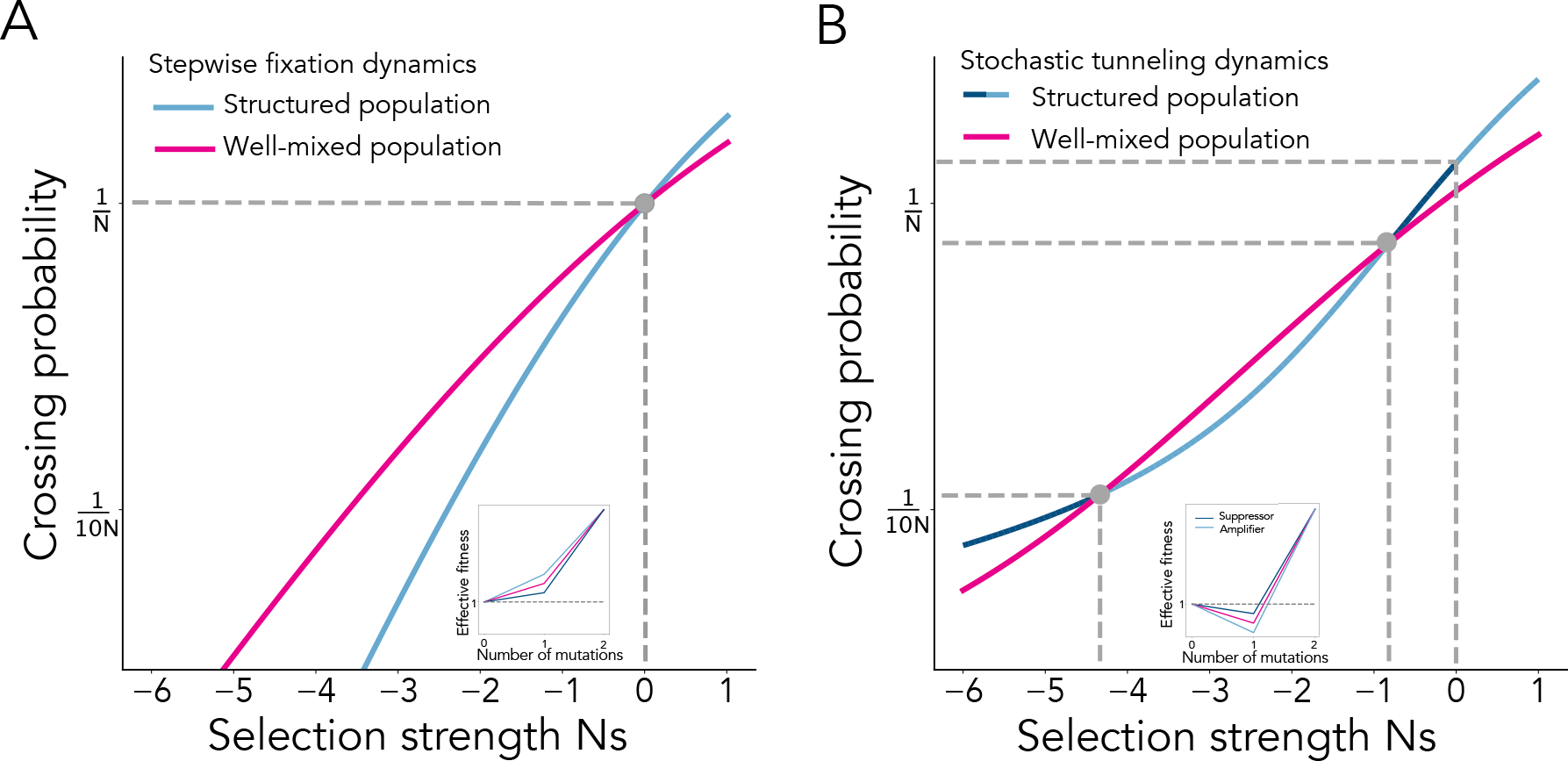
Complex population spatial structure reshapes the effective fitness landscape. Here, the color of the lines represents amplification (light blue) or suppression (dark blue) of selection. The population size is 1000. **A. Stepwise fixation dynamics.** We compare fixation probabilities of the second mutation in a structured population (*α* = 1.5, *λ* = 4) and a well-mixed population, in the limit of weak mutation (*µ* 0). An amplifier of selection increases the effective fitness of the intermediate mutant making the fitness landscape more rugged. In contrast, a suppressor of selection would decrease the effective fitness of the intermediate mutant, smoothing out the fitness landscape. In this regime, the dynamics can be predicted using single-mutation results. **B. Dynamics under stochastic tunneling.** We compare fixation probabilities of the second mutation in a structured population (*α* = 1.5, *λ* = 4) and a well-mixed population under stochastic tunneling (*µ* = 10*^−^*^7^). In this regime, population structure crosses the wellmixed line twice, for negative selection strengths, and keeps switching from amplifying to suppressing fitness landscape crossing and fixation of the second mutant. This regime is harder to understand and predict using single mutation results.

In contrast, a stochastic tunneling regime allows for the fixation of the second mutation without the intermediate mutation first sweeping to fixation. This causes the evolutionary dynamics to deviate significantly from the case of step-wise fixation. Intuitively, this happens because landscape crossing is fundamentally shaped by both the probability and the time it takes for the first mutant to spread in the population and create opportunities for the appearance of the second mutant. This interplay between how the spatial structure changes the fixation probability and the fixation time of the intermediate mutant leads to rich landscape crossing dynamics that cannot be predicted or interpreted through the simpler lens of single mutation fixation probabilities.

In previous work, we have shown that, for single mutations, the evolutionary role of complex spatial structure can be quantified by analyzing two essential network properties: the network amplification factor, which shapes probabilities of fixation compared to well-mixed populations (Lieberman et al., 2005; Kuo et al., 2021) and the network acceleration factor, which shapes the time to fixation for new mutations in the population (Kuo and Carja, 2021). The network amplification factor *α* rescales *s* into an effective selective coefficient, *s_e_* = *αs*. Similarly, the network acceleration factor, *λ*, shapes times to fixation by essentially changing the effective population size to *N_e_*= *λN* (Kuo and Carja, 2021).

We find that, under a stochastic tunneling regime, the network amplification factor is no longer predictive of the new effective selection coefficient and the intercept between the crossing probability of the structured population and that of the well-mixed case can shift from neutrality (**Figure 2B**). For example, for the amplifier of selection in **Figure 2B**, there exists a region between the new intercept and neutrality, where the population structure behaves as a suppressor (dark blue portions of the line). In addition, there exists an additional intercept, where the population structure acts as a suppressor on the left and an amplifier on the right. This example highlights that single mutation fixation probabilities can lead to erroneous inferences of multi-mutational landscape crossing dynamics: there exist regions of the parameter space where an amplifier of selection for a single mutation acts instead as a suppressor of selection on the landscape.

We start by intuitively describing the roles of the two network parameters in shaping fitness landscape crossing and introducing the main ideas of our full analytic approach.

**The role of the network acceleration factor.** Let us assume a network that only affects time to fixation and not the probability of fixation (*α* = 1) for a single mutant. There are two independent evolutionary scenarios that can lead to the acquisition of the second mutant.

The first scenario occurs when the intermediate mutant lineage is poised for extinction and the second mutation appears in the population before the lineage goes extinct. In a well-mixed population, the lineage reaches a size *T*, smaller than the establishment threshold of 1*/s*, in *T* generations and then drifts to extinction in another *T* generations, with probability 1*/T* . The overall shape of such a trajectory is shown in **Figure 3A** and the total number of mutational opportunities to acquire the second mutation depends on the area under the trajectory, *W* (*T*) = *T* ^2^ (Weissman et al., 2009). The probability that a second mutation arises on the background of this mutant lineage is the expectation of the probability that a second mutation occurs given trajectory *T*, Φ(*W* (*T*)), over all possible trajectories *T*,

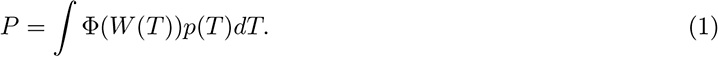

**Figure 3:**
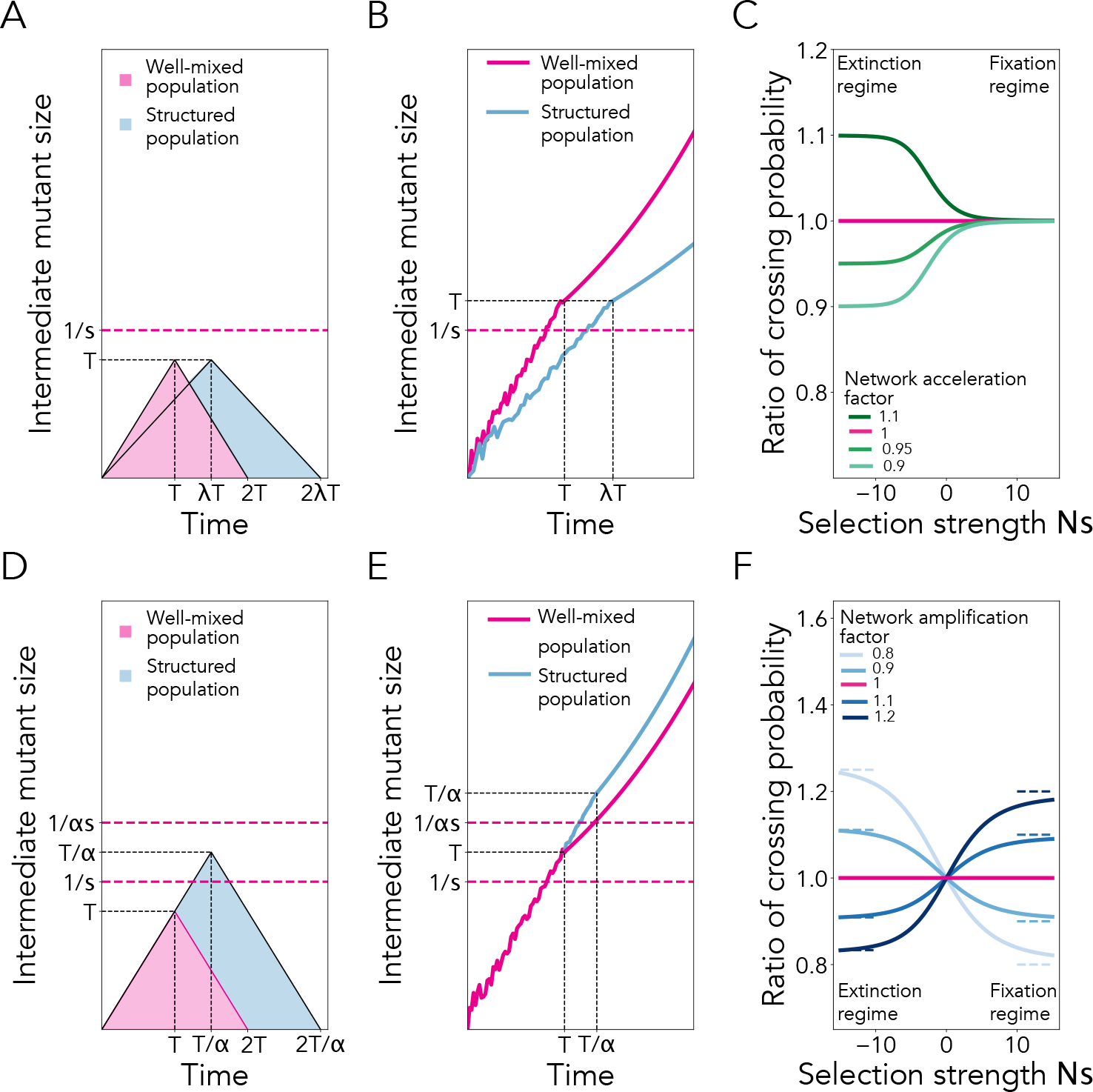
The fate of the intermediate mutant provides intuition of how spatial structure reshapes crossing dynamics. Top panels show a population structure with amplification equal to one. **A.** Typical trajectory of the intermediate mutant frequency, given that the intermediate goes extinct. Population structures that increase extinction time lead to an increased probability of acquiring the second mutation. **B.** Typical trajectory of the intermediate mutant frequency, given that the intermediate mutant fixes. **C.** Ratio of crossing probability between the network-structured population and the well-mixed model. Bottom panels show a population structure with acceleration equal to one. **D.** Typical trajectory of the intermediate mutant frequency, given the intermediate mutant goes extinct. A suppressor increases the line of transition between the stochastic and the deterministic dynamics, allowing the mutant to drift for longer and to a higher frequency. The structure would, therefore, increase the probability of acquiring the second mutation. **E.** Typical trajectory of the intermediate mutant frequency, given the intermediate fixes. **F.** Ratio of crossing probability between the network-structured population and the well-mixed model. The dashed line represents approximation in equation 4: on the left, the line is given by 1/*α* and on the right, the line is given by *α*.

When the magnitude of *s* is strong enough (*|s| > ^√^µ*), the rate at which double-mutants are produced is dominated by rare lucky intermediate mutant lineages that survived for *T ∼* 1*/s* (Weissman et al., 2009). Therefore, the probability of landscape crossing under this scenario is

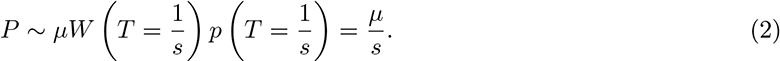

A network-structured population with acceleration factor *λ* effectively stretches the time the lineage reaches a size of *T* to *λT* generations, and also the time to extinction to 2*λT* generations. The area under the trajectory thus becomes *λT* ^2^ and, using equation (2), the rate of landscape crossing is changed by a factor of *λ,* to *µλ/s*.

The second scenario occurs when the intermediate mutant lineage would reach fixation on its own. In this scenario, the trajectory behaves stochastically below the establishment threshold, 1*/s*. As soon as the mutant frequency crosses this threshold the population dynamics becomes essentially deterministic (**Figure 3B**). At any point along the trajectory, the second mutation can occur and carry the mutant lineage to fixation. Since this scenario is conditioned on the first mutant fixing, the population is thus guaranteed to eventually acquire the second mutation. Therefore, in this regime, the network structure does not change the probability of crossing the fitness valley.

The total probability of crossing the fitness landscape is the sum of the probabilities of acquiring the second mutation under the two independent evolutionary processes (**Figure 3C**). When the first mutant is strongly deleterious, the population depends on the second mutation appearing in time to cross the fitness landscape and the acceleration factor of the network changes the rate of fitness valley crossing by a factor of *λ*. Otherwise, the probability of stochastic tunneling remains unchanged by the network structure.

### The role of the network amplification factor

Let us now assume a network structure that changes the fixation probability of a new mutant, but not the time to fixation (*λ* = 1) (**Figures 3D** and **E**).

For a network with amplification factor *α*, the mutant experiences an effective selection strength of *αs*.

Therefore, if the intermediate mutant lineage is on its way to extinction, with probability 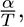, the mutant reaches an intermediate mutant size of 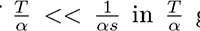 generations and goes to extinction in another 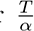 generations. The network amplification factor does not change the expected trajectory that reaches size 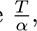, but instead changes the limit on the size that an intermediate mutant trajectory can reach, from a maximum of 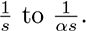. Using equation (1), the rate of landscape crossing under this scenario becomes *µ/αs*. In contrast, if the intermediate mutant linage is destined to reach fixation, the crossing probability is the same as the probability of fixation of the first mutant with effective fitness *αs*, since the population is guaranteed to acquire the second mutation.

In this regime, the network structure does not change the probability of crossing the fitness valley, beyond modifying the effective selective coefficient of the intermediate mutant.

Considering the two scenarios together, when the first mutant is strongly deleterious, the population depends on the first scenario to cross the fitness landscape, and the rate of landscape crossing is changed by a factor of 1*/α*. In the limit where the first mutant is strongly beneficial, fixation of the intermediate becomes the dominant process, and the crossing probability is approximately *αs* (**Figure 3F**).

### The general analytic approximation

Using the intuition above, we approximate the landscape crossing probabilities for an arbitrary network structure with amplification parameter *α* and acceleration parameter *λ*, in the limit of large population size (for the full description of our approach see the **Supplementary Material**, in particular equations (36)-(44)). We approximate the crossing probability to be

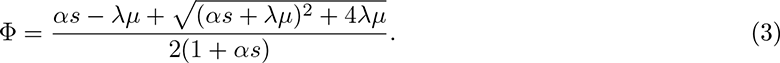

Under weak mutation, series expansion in *µ* (see **Supplementary Material** equations (47)-(51) for the full description) reduces equation (3) to

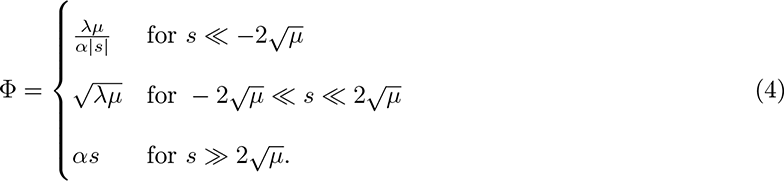

In the deleterious intermediate mutation regime, *s « −*2*^√^µ,* the mutant lineage goes to extinction with high probability if the second mutation does not appear. Equation (4) recaptures the crossing probability of *λµ/|s|* when *α* = 1, and *µ/α|s|* when *λ* = 1. A network structure, with amplification parameter *α* and acceleration parameter *λ* changes the crossing probability of the well-mixed population, *µ/|s|*, by *λ/α*. Therefore, an amplifier, as defined in the single mutant case, can promote the fixation of a mutant linage with deleterious intermediates, as long as the acceleration parameter is greater than the amplification parameter. This is in stark contrast to the case of sequential fixation, where an amplifier always reduces the acquisition probability of the second mutation, with a deleterious intermediate mutant. With neutral intermediate mutants, *−*2*^√^µ « s «* 2*^√^µ,* selection on these mutants has a negligible effect on the crossing probability: equation (4) only depends on *λ* and not *α.* In the case of a beneficial intermediate mutation, 2*^√^µ » s,* the population crosses the fitness landscape through intermediate mutants that are destined to fix. Since *λ* only affects the probability of acquiring the second mutation before fixation of the first, the crossing probability only depends on *α.* Here, the crossing dynamics reduces to the sequential mutation case.

When crossing a fitness landscape, the spatial structure of the population can still act as an overall suppressor or amplifier, but only in a small region of the parameter space where the acceleration factor is exactly one. Beyond this knife-edge case, the general dynamics become more complicated, with the existence of piece-wise acceleration or suppression and we observe seven additional categories of graphs as the acceleration factor moves away from one (**Figure 4**).

**Figure 4:**
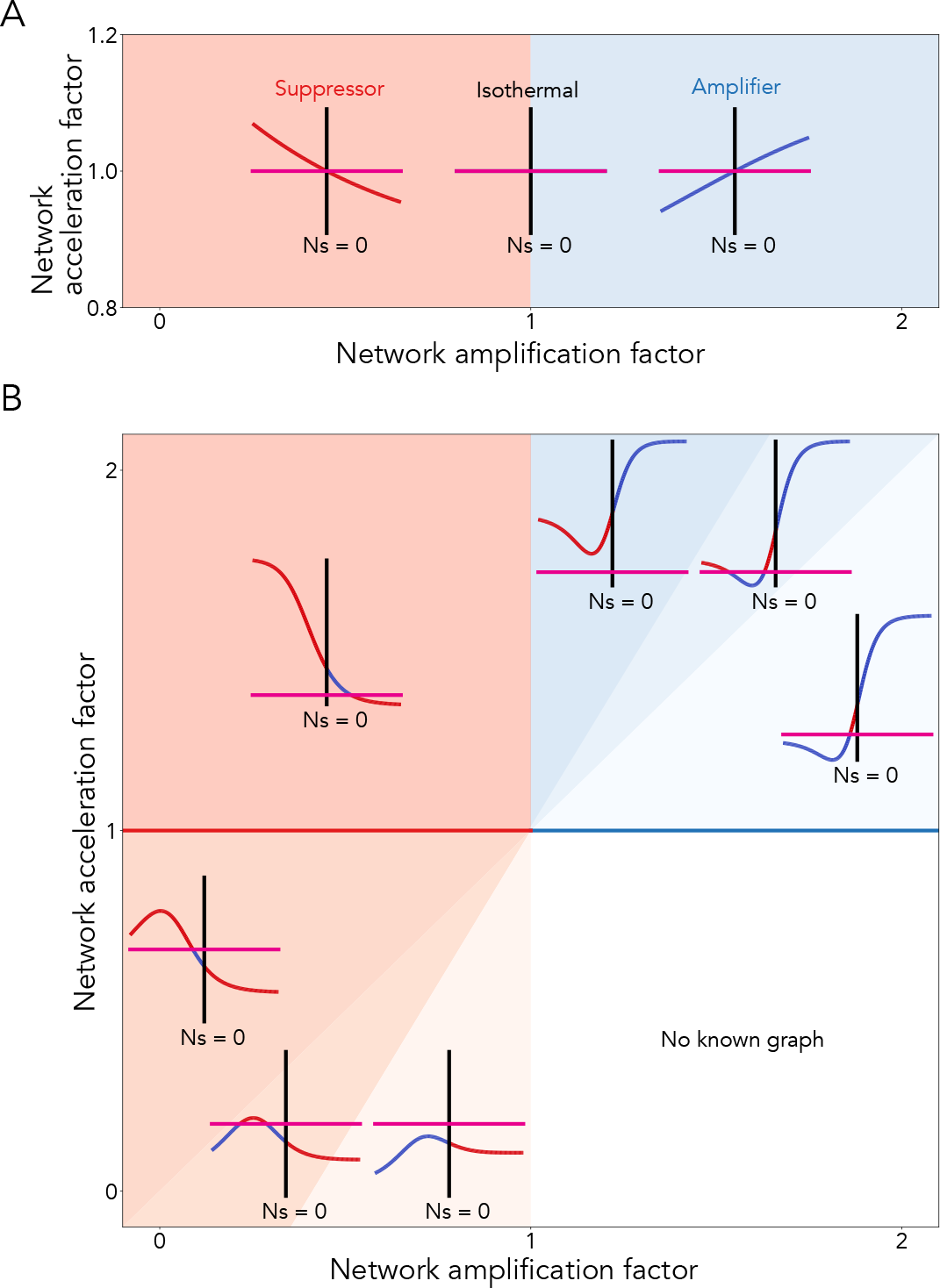
**A unified view of how different categories of graph structures reshape fitness landscape crossing dynamics. A**. The stepwise fixation regime (*µ* 0). **B**. We analytically calculate the crossing probability based on the amplification *α* and acceleration *λ*, for graphs of size 1000. The y-axis on the inlet is the ratio of crossing probability, and the x-axis is the selection strength. The color of the line represents amplification of selection: blue for increased selection pressure, and red for decreased selection pressure. The pink line is the ratio of crossing probability for the well-mixed model. Here, the mutation rate is set to 10*^−^*^6^.

These additional categories can be understood using the intuition developed in **Figure 3** and equation (4). A network structure with *α <* 1 and *λ >* 1 (*λ/α >* 1) acts as a suppressor, increasing the crossing rate for a deleterious intermediate (the upper left region of **Figure 4B**). For a neutral mutation, the crossing probability depends on *^√^λ.* Therefore, for *λ >* 1, the ratio of crossing probability crosses neutrality above the well-mixed line and the population structure acts as an amplifier, for weakly beneficial mutation. As the strength of selection increases, the crossing probability depends on 1*/α.* Since *α <* 1, the population structure acts as a suppressor, decreasing the the rate of valley crossing. As *λ* decreases, and 1*/α <* The overall dynamics*_√_*i<u>n</u> the inlet can be explained through equation 4: the line approaches *λ/α* on the left, crosses neutrality at *λ*, and approaches *α* on the right. 1, the the population structure becomes an amplifier for weakly deleterious mutations. As *λ* decreases even further, *λ/α* is no longer larger than one. The population structure starts behaving like an amplifier for deleterious mutations, crosses neutrality below well-mixed, and behaves as a suppressor for beneficial mutations. There is a sub-category of behavior where the ratio of crossing probabilities is higher than one for an intermediate deleterious mutation. This is not predicted by equation (3), since the approximation assumes a large enough population size, such that deleterious intermediates fix with negligible probability. For smaller population sizes however, the deleterious intermediate can be carried to fixation just through drift. A population structure with an amplification factor less than one can thus increase the fixation probability of this intermediate, leading to a higher crossing rate.

In the regime where *α >* 1, a population structure with *λ »* 1*/α* will increase crossing probability compared to a well-mixed population, regardless of the fitness of the intermediate. Population structures with multiple transitions between amplification and suppression occur when *λ →* 1*/α*, as *λ* decreases towards one. No graphs exist in the bottom left corner of the parameter space, since networks that are amplifiers have longer fixation time than well-mixed populations (Tkadlec et al., 2019).

### The roles of other network parameters

While the network amplification and acceleration factors play a unifying role in shaping evolutionary dynamics on the network, we also analyze how other network properties, in particular network connectivity and node heterogeneity, shape the fitness landscape crossing probability. We use preferential attachment networks, which allow for easy independent tuning of the number of edges and the shape of the degree distribution. For a deleterious intermediate mutant (here *Ns* = *−*1), we show that the crossing probability decreases with increasing mean degree of the network and increases with degree heterogeneity when the mean degree is low (**Figure 5A**). This is a marked deviation from previous results. Classic stepwise fixation accounts for around 95% of a total successful crossing. Since the second mutant is assumed to fix immediately upon introduction, the crossing probability under stepwise fixation depends solely on the fixation probability of the intermediate mutant. Preferential attachment graphs with low mean and high variance in degree are known to be strong amplifiers of selection, reducing the probability of fixation of a deleterious mutant. One would expect these network structures to have a lower crossing probability than that of the well-mixed population, the opposite of what we observe in **Figure 5A**.

**Figure 5:**
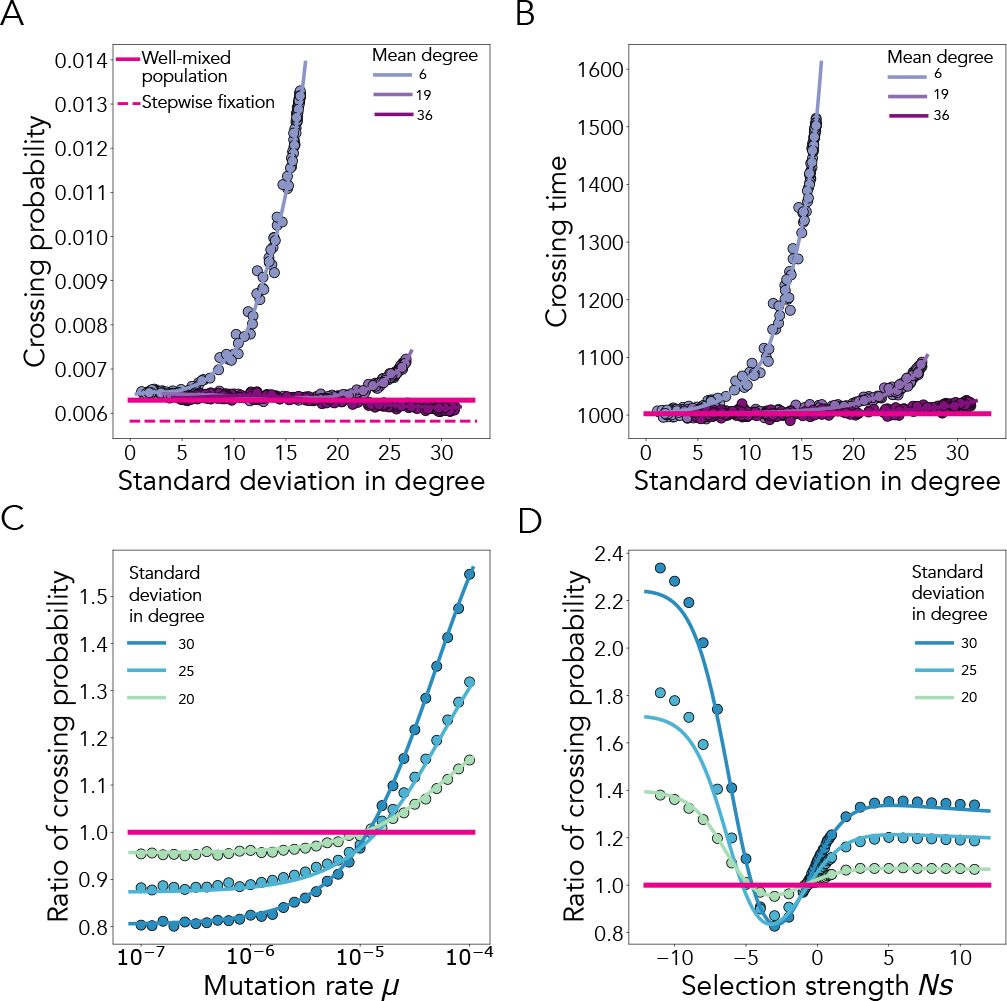
Complex population structure can promote crossing of fitness landscapes, even if the intermediate mutation is deleterious. The dots represent ensemble averages across 10^7^ replicate Monte Carlo simulations, while the lines represent our analytical approximations. **A.** Here, the color indicates the mean degree of the network, as in the legend. For dots of the same color, the mean degree of the network is held constant, as we vary the variance of the degree distribution. Population size N = 100, *µ* = 10*^−^*^5^ and s = -0.01, such that Ns = -1. **B.** The crossing time across networks with different variance of the degree distribution. Here, N = 100, *µ* = 10*^−^*^5^ and s = -0.01, such that Ns = -1. **C.** The crossing probability, as a function of mutation rate. Here, N = 100, mean degree is 19 and s = -0.01. **D.** The crossing probability, as a function of selective pressure. Here, N = 100, *µ* = 10*^−^*^5^ and mean degree is 19.

Since we are considering evolutionary scenarios where the mutation rate is high or the population size is large, the rate of evolutionary dynamics in the population depends not only on the probability of crossing, but also on the expected time to cross. For the same mutational landscape as **Figure 5A**, the crossing time decreases with increasing mean degree and increases with increasing network node heterogeneity (**Figure 5B**, see **Supplementary Material** for the analytic derivation). There is a positive correlation between crossing probability and time, similar to the single mutation case, which highlights the tradeoff between probability and time when designing population structures for evolutionary optimization (Möller et al., 2019; Kuo and Carja, 2021).

We show the shift of dependence on stepwise fixation to stochastic tunneling, as the mutation rate increases, in **Figure 5C**. We quantify the difference in the likelihood of crossing the fitness landscape by dividing the crossing probability of the structured population by that found in the well-mixed model. When the mutation rate is low, the second mutant fixes solely through fixation of the first mutant. The ratio of the crossing probability is then simply the ratio of the fixation probability of the first mutant. Since the intermediate mutant is deleterious, the results here are consistent with the fact that preferential attachment graphs are amplifiers for selection for a single mutant. As the mutation rate increases, stochastic tunneling starts to become a dominant contributor to the crossing probability and the networks begin to act as a suppressor of the first mutation, thereby increasing the crossing probability compared to the well-mixed model.

We show that our approximation holds across a range of mutational landscapes, ranging from fitness valleys to fitness plateaus and fitness hills. There is a nonlinear behavior in the relative likelihood of landscape crossing as a function of selection strength (**Figure 5D**). In the strongly deleterious valley crossing regime, we see the largest deviation from the known theory of single mutation dynamics. This is caused by the fact that strongly deleterious mutations are unlikely to fix, thus the successful crossing of the landscape depends solely on stochastic tunneling. As intermediate fitness increases, fixation of the first mutant becomes increasingly likely, adding a larger contribution to the crossing of the fitness landscape, and the evolutionary dynamics converges to that of the single mutation dynamics.

### Evolutionary landscape crossing across different network families

Our analytical results provide a unifying way of explaining complex multi-mutational dynamics by distilling properties of complex network organization through two single parameters: the acceleration and amplification factors. We can use this framework to compute the rate of fitness landscape crossing across all families of networks.

For a given landscape ruggedness, we can design networks with the right ratio of amplification and acceleration to optimize for the rate of valley crossing. For example, for a deleterious intermediate mutant, since the crossing probability depends on *λ/α*, the crossing probability increases with increasing acceleration factor and decreasing amplification factor (**Figure 6A**). With an elevated mutation rate, the crossing time becomes a major contributor to crossing rate, alongside the crossing probability. Since the intermediate mutant lineage must wait for *∼ λ/α* for the second mutation to occur, the crossing time must also scale with *α/λ* (top of **Figure 6B**). As the acceleration factor decreases, tunneling is unlikely, since the mutant lineage goes extinct before the second mutation can occur. The population has to rely on sequential fixation to cross the fitness valley. Therefore, in a population with a low acceleration factor, the fixation time is negligible compared to the expected time for mutation, and the crossing time depends on 1*/Nµ*. Crossing by stochastic tunneling has expected wait time of *∼ λ/α,* which is much lower than 1*/Nµ*. A population structure with a higher acceleration factor, where tunneling becomes likely again, would cross the fitness valley at a faster rate compared to a population with a lower acceleration factor.

**Figure 6:**
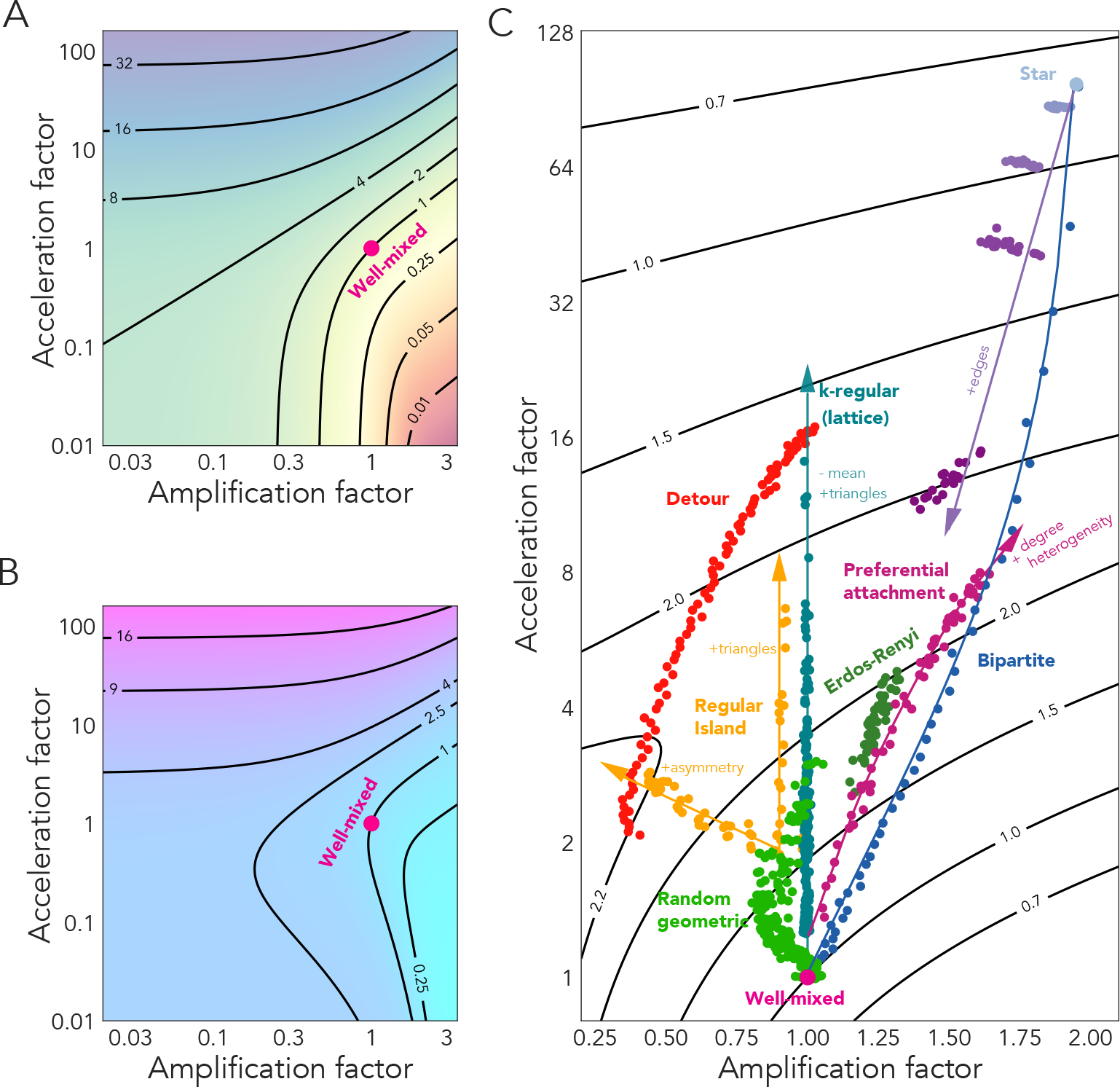
The mutational landscape determines which network structure maximizes the rate of evolution. We plot the rate of fitness valley crossing, as function of the amplification and acceleration factor for graphs of size 100. The mutation rate is set to 10*^−^*^4^. For the intermediate mutant, *Ns* = 5. **A.** The color represents the ratio of crossing probability compared to the well-mixed population. **B.** The color represents the ratio of crossing time, compared to the well-mixed model. **C.** The black lines are contour lines of the ratio of crossing rate and the dots represent individual networks of the families depicted.

Using the crossing probability Φ and crossing time *τ,* we approximate the effective rate of crossing a fitness landscape by 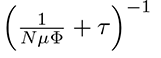 (Frean et al., 2013). Using this approximation for the rate of evolution, we compare the main families of networks in **Figure 6**C. The star graph has the highest crossing probability due to its ability to prolong the persistence of mutants, despite being an amplifier. For the same reason, the mutants also take a long time to reach a larger size for tunneling to occur, leading to a long crossing time. These two contributors to the rate of evolution put together lead to an overall lower rate of crossing for the star graphs. The well-mixed population, on the other hand, has a low acceleration factor, allowing for the intermediate mutant frequency to change on a faster time period. As a result, the well-mixed population has a fast crossing time, but a low crossing probability. This also makes the well-mixed population sub-optimal for crossing this fitness valley. The asymmetric regular islands and detour graphs have the highest crossing rate out of all graphs we explored. These graphs are suppressors for single mutations and have moderate acceleration factors that give the right balance of crossing probability and crossing time. Another graph family of interest is the bipartite graph. These graphs have the lowest fixation time for a given fixation probability of a single mutation. However, they span a wide range of crossing rates, none of which are optimal. This reinforces our previous result that a network that is Pareto optimal for a single mutation in terms of fixation time and probability does not guarantee the best performance on rugged fitness landscapes. The best-performing graph is inherently dependent on the underlying mutation landscape.

### Rates of evolution to leukemia initiation in the bone marrow

We apply this model to study the multi-hit process of leukemia initiation in the bone marrow as a function of the spatial organization of the stem cell niches, mutation rates and distribution of mutational effects. Mathematical models based on known rates of cancer incidence suggest that combinations of more than two oncogenic mutations are required for carcinogenesis (Knudson Jr, 1971). These mutations often involve tumor suppressor genes, which require knock-out mutations in both copies of the chromosome (**Figure 7A**). Here we assume there exists a set of cancer driver mutations with different effects on cell fitness and the acquisition of any pair of genes in the set leads to leukemia initiation. In line with our model, the cancer-initiating mutant is assumed to have an extremely high somatic fitness and to fix upon introduction. We assume a negative gamma distribution for the distribution of mutational effects (**Figure 7B**), shifted to the right to account for beneficial mutations (Piganeau and Eyre-Walker, 2003). For a range of biologically relevant mutation rates, and using previous estimates of amplification and acceleration factor for the network structures of the stem cell niches of the bone marrow (Kuo et al., 2021), we use our analytic results to compute the total rate of neoplasm initiation,

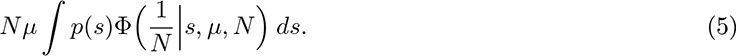

**Figure 7:**
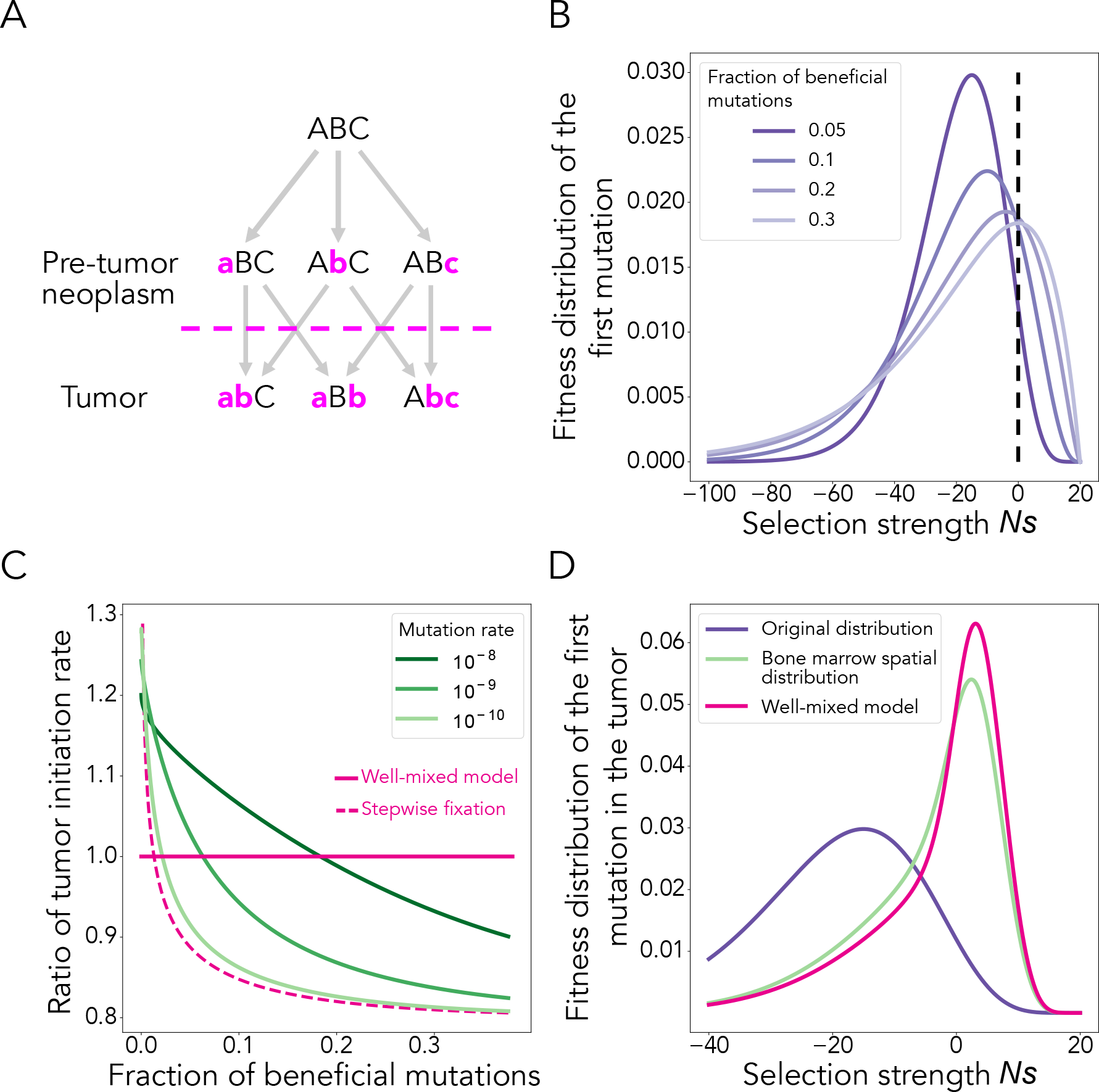
A 2-hit model for tumor initiation predicts bone marrow architectures to suppress mutation accumulation, if beneficial mutations make up at least a small portion of driver mutations. A. Illustration of a 2-hit model. In this example, there are 3 genes where any combination of two mutations leads to cancer. The letters represent a genotype and the arrows represents mutation paths. **B.** We assume the intermediate mutations to have different fitness effects, as represented by a shifted negative gamma distribution. **C.** The ratio of cancer initiation rate between the bone marrow tissue cellular architecture and the well-mixed population, as a function of the shape of the fitness distribution and the mutation rate. Here, the gamma distribution has a fixed mean selection strength of *Ns* = 20. The population size is 50000. **D.** The difference in the posterior fitness distribution of first mutant, between the well-mixed and the network structure representative of the stem cell niches of the bone marrow. Here, the gamma distribution has a fixed mean selection strength of *Ns* = 20 and a standard deviation of 75. The mutation rate is 10*^−^*^9^.

We have previously shown that the networks of the stem cell niches of the bone marrow are suppressors of selection (Kuo et al., 2021), therefore the rate of crossing fitness hills should be decreased and the rate of crossing fitness valleys should be increased. We find that the change in the rate of cancer initiation, compared to a well-mixed population, depends heavily on the assumed fraction of incoming beneficial mutations into the population (**Figure 7C**). At smaller, native mutation rates, the bone marrow mostly suppresses the rate of leukemia initiation, but at higher mutation rates, under environmental carcinogens, for example, the point of crossing the well-mixed regime is shifted to the right, and, over a wider space of parameters, the tissue increases the rate of cancer initiation over the well-mixed population.(**Figure 7C**)

The population structure of the bone marrow also better tolerates intermediates with deleterious effects, than a well-mixed cellular population. **Figure 7D** shows the distribution of fitness effects for intermediate mutations entering the population, alongside the fitness distribution of the intermediates found after fixation of the second mutation. When studying cellular populations that successfully crossed the somatic evolutionary landscape, such as those in tumor samples, the distribution of fitness effects observed will be different as a function of the spatial cellular architecture. In **Figure 7D**, the fitness distribution of mutations entering the population is mostly deleterious, but the fitness distributions of intermediate mutations that lead to fixation of the lineage, for both well-mixed and bone marrow populations, are biased towards beneficial mutations. This is intuitive, since the beneficial intermediate can reach fixation on its own and wait for the second mutation to appear, while the deleterious intermediate must rely on stochastic tunneling. As a result, the advantageous intermediates are much more likely to be observed when looking at the distribution of fitness for the populations that have crossed the fitness landscape. This distribution is skewed to the left for the bone marrow architecture, implying that the bone marrow favors deleterious intermediates more than a well-mixed population. Our results therefore showcase how tissue architectures can influence paths taken by the population across fitness landscapes, and this is reflected in the observed posterior mutational fitness distribution.

## Discussion

Here we develop a unifying theory of evolutionary dynamics on graph-structured populations beyond single mutational steps. We show that network-structured populations can increase the crossing rate across a rugged fitness landscape, compared to well-mixed populations, by extending the time at which the deleterious and neutral intermediate go extinct. This is in agreement with previous lattice and deme-based models which show that these regular spatial structures can permit intermediate mutants to persist, until the beneficial final mutation occurs (Komarova, 2014; Bitbol and Schwab, 2014). Our results extend these initial findings and analyze non-symmetric spatial structures of controllable complexity. In stark contrast with the symmetric models, these population structures also change the probability that the deleterious and neutral intermediates go extinct and this can produce a much wider range of evolutionary outcome.

Using simulation and analytic approximations, we show that these complex evolutionary dynamics can be intuitively explained by analyzing how two main network properties, the amplification and acceleration factors, change the expected fate of the intermediate mutant in the population. Importantly, this intuition can inform on the design of population topologies that optimize rates of crossing fitness landscapes: 1) network-structured populations with a high ratio of acceleration to amplification factor cross fitness valleys and plateaus more effectively and 2) network populations with high amplification factor cross fitness hills more effectively. The first observation can be understood and interpreted using Wright’s shifting balance theory, since through either decreased effective selection or increasing hindrance to gene flow, an increase in the relative force of drift leads to an increased window in which the final beneficial mutation can rescue the mutant lineage (Wright et al., 1932).

Our results generalize to more complex mutational landscapes with more than two mutational steps and the model can be easily extended to accommodate a *K*-step mutational process, even when the end mutant is not guaranteed to fix. This can be done by sequentially transforming the *K*-step process into a two-step process, where the probability of reaching the *K*-th mutant from the first mutant is treated as the probability of fixation of the second mutant in a transformed process. Iteratively repeating this process allows us to find the crossing probability for the entire *K*-step process. Using previous results (Weissman et al., 2009), we can infer a simple expression for the crossing probability:

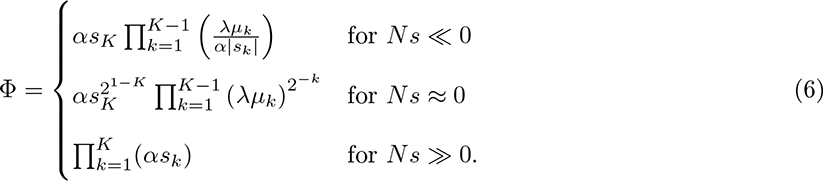

The two-step process allows us to showcase the complex evolutionary scenarios enables by the addition of just one mutational step, while also being simple enough to draw direct comparisons with the case of a single mutation and the amplifier / suppressor framework. However, as long as the fitness landscape is consistent, meaning the landscape consists of only hills, plateaus, or valleys and does not exhibit complex ruggedness, then a population structure that promotes landscape crossing for the two-step process, will act similarly for the K-step process. On the other hand, on a rugged fitness landscape, with arbitrary signs and strength of selection across the landscape, the amplifier / suppressor descriptive framework can become illusive. Further exploration is needed to link existing measures of landscape ruggedness with measures of network topology for building a unifying theory for complex fitness landscapes.

We categorize the different evolutionary scenarios through analysis of the transitions between when the graph acts as an amplifier, versus as a suppressor. We find that most network structures are rarely pure amplifiers or suppressors in the two-step process, instead transitioning from one to another. Two categories of the observed dynamics are of particular interest: amplifier to suppressor and, separately, suppressor to amplifier. A network that falls in the amplifier to suppressor regime will decrease the landscape crossing probability compared to a well-mixed model, regardless of selection strength of the intermediate mutant.

A graph with such properties in the case of single mutation was coined as a reducer, and the ring graph under the death-Birth process is a well-known example (Allen et al., 2020; Hindersin et al., 2016b). A network that falls in the suppressor to amplifier regime will always increase the crossing probability for all intermediate mutations, whether beneficial or deleterious. We call this type of graph an enhancer of landscape crossing. Regular graphs that only change single mutation fixation time are examples of this (**Figure S2**). To our knowledge, no network structure under any update process, whether Birth-death or death-Birth, was previously found to be an enhancer of fixation for the single mutation case. If the ruggedness of the landscape is known, networks that specialize in a specific landscape can overall perform better than enhancers of crossing and our analytic results can be easily used to design and select the ideal network topology. In cases where the exact shape of the optimization landscape is unknown, often observed in reinforcement learning problem where the agent needs to sample from the environment to get an estimate of local information, and enhancer graphs become the obvious best choice for their ability to speed up learning across different landscapes (Williams, 1992; Peters and Schaal, 2008; Schulman et al., 2015).

One key limitation of our analysis is the focus on low selection strength for the intermediate mutant. In this selection regime, the amplifier / suppressor framework, with constant strength of amplification with respect to selection is a valid first-order approximation (McAvoy and Allen, 2021). Under a broader range of selection, there is a wide range of graphs that are piece-wise or transient amplifiers / suppressors, with the graphs having selection-dependent amplification factors (Voorhees, 2013; Voorhees and Murray, 2013; Allen et al., 2020). Finer categorizations of amplifiers also exist when considering different mutant initializations, where the mutant does not appear uniformly in the population (Allen et al., 2021). Extending the analytic treatment of crossing dynamics to these selection regimes is challenging, since exact calculation grows exponentially with population size Hindersin et al. (2016a) and stochastic tunneling often occurs in large populations.

Our model can be applied to understand the evolutionary trajectory of cellular systems with complex spatial architectures, and we use an existing dataset to study how the spatial organization of the stem cell niches in the bone marrow shapes rates of leukemia initiation. We find that the relative rate towards neoplasm initiation in this spatial cellular architecture depends critically on the mutation rate, as well as the proportion of beneficial mutations coming into the population. We show that a structure that suppresses cancer initiation can instead transition into one promoting cancer initiation, depending on the distribution of mutational effects. This result has wide clinical implications. A previous analysis of nearly 50,000 healthy individuals showed that most somatic mutations confer a fitness advantage (Watson et al., 2020).

When driver mutations with fitness advantages are prevalent, the bone marrow architecture suppresses accumulation of driver mutations. We also show that under exposure to carcinogens (increased rate of mutation), the architecture of the bone marrow increases rates of neoplasm initiation. In fact, increasing mutation rate is a known hallmark of cancer (Nowak et al., 2002; Pino and Chung, 2010; Tomlinson et al., 1996). One can therefore hypothesize that tumors take advantage of a normal tissue structure, evolved to suppress mutation accumulation, and, by increasing the mutation rate in the population, effectively turn the same tissue structure into a promoter of further mutation accumulation to combat immune response and clinical intervention.

## Acknowledgments

We gratefully acknowledge support from the NIH National Institute of General Medical Sciences (award no. R35GM147445), the United States-Israel Binational Science Foundation (award no. 2019266) and from the NIH T32 training grant (no. T32 EB009403). This research was done using resources provided by the Open Science Grid, which is supported by the National Science Foundation award 1148698, and the U.S. Department of Energy’s Office of Science.

