## Supplementary Material for "Evolutionary graph theory on rugged fitness landscapes"

**2 Landscape crossing results on various graphs families** **15**

**3 Application to the bone marrow stem cell population architectures. Rates of neoplasm initiation.** **21**

### 1 The analytic approach

In this section, we describe our analytical approach to the problem of fitness landscape crossing for graph structured populations. We show how we derive the approximations used in the main text, for the two quantities of interest (as proxies for reaching the fitness peak): the probability that a second mutation arises in the population, and the expected time for that to occur. In what follows, we will use the diffusion approximation. Direct application of the diffusion approximation however is difficult due to the complex dynamics that can occur on a graph. For that reason, we instead split the analysis into two parts. In the first part, we begin by describing the deterministic population dynamics and we identify the neutral equilibrium of the mutant in the population. This neutral equilibrium describes the distribution of the mutant over the nodes of graph, assuming zero stochastic fluctuations. In the second part, we use this deterministic mutant distribution, apply the diffusion approximation and analyze the stochastic dynamics of the system.

#### 1.1 The deterministic dynamics

We assume a two-step mutation process in a population of fixed finite size $N$. We denote the mutation rate by $\mu$ and we ignore back mutations. We assume two different types of individuals in the population: a wildtype $A$ and the mutant lineage $a$. Here $a$ denotes both the intermediate mutant lineage and the two-mutant lineage, since we assume the final mutant fixes immediately upon appearance in the population. The wild-type has reproductive fitness of 1, while the intermediate mutant has a fitness of $(1+s)$. We define $n_a$ and $p_a$ as the number and frequency of mutant individuals in the population. Similarly, $n_A$ and $p_A$ denote number and frequency of wildtype in the population.

Spatial structure is represented by a graph, where $n_i$ and $p_i$ denote the number and frequency of node of degree $i$, and $n_{ij}$ and $p_{ij}$ denote the number and frequency of edges that connect node of degree $i$ to nodes of degree $j$. Let $n_{i,a}$ be the number of mutants that occupy nodes of degree $i$ and $n_{ij,Aa}$ be the number of edges that connect nodes $A_i$ and $a_j$, nodes of degree $i$ containing the wildtype with nodes of degree $j$ containing the mutant lineage. Similarly, we define $n_{ij,AA}$ and $n_{ij,aa}$. We also define frequencies $p_{i,a}$ as $n_{i,a}$ divided by the total population size and $p_{ij,Aa}$ as $n_{ij,Aa}$ divided by the total number of edges $\sum n_{ij}$.

We first derive the equations governing the mutant frequency dynamics on nodes of degree $i$, $p_{i,a}$. We show that $p_{i,a}$ depend on the frequencies of edge types in the graph, $p_{ij,AA}$, $p_{ij,aa}$ and $p_{ij,Aa}$. By deriving the edge frequencies, we show that the edge dynamics further depend on higher order mutant configurations on the graph, such as $p_{ijk,AAa}$. To simplify the mathematical analysis, we can approximate these higher order mutant configurations using just the frequencies of the mutant on nodes of degree $i$ and the frequencies of edges connecting mutant and wildtype nodes. We use these approximations to derive the equilibrium mutant frequencies $p_{i,a}$. These equilibrium frequencies are critical for then deriving and understanding the mutant stochastic dynamics on the graph.

##### 1.1.1 Mutant frequency dynamics on nodes of degree $i$

The probability that a mutant occupying a node of degree $j$ replaces a wildtype occupying a node of degree $i$ is the probability of selecting a node $a_j$ to reproduce and a neighboring node $A_i$ to die. This probability is given by

$$\begin{aligned} P(a_j \to A_i) &= (1-\mu)\frac{1+s}{Nw}n_{j,a}\frac{n_{ij,Aa}}{jn_{j,a}} \\ &= (1-\mu)\frac{1+s}{Nwj}n_{ij,Aa}. \end{aligned} \tag{1}$$

Here, $w = p_A + p_a(1+s)$ represents the mean fitness of the population. Using this probability, we calculate the expected change of numbers of different node types, in one update as

$$\begin{aligned} a_j \to A_i: \quad \Delta n_{i,a} = +1 \quad \mathbb{E}[\Delta n_{i,a}|a_j \to A_i] &= (1-\mu)\frac{1+s}{Nwj}n_{ij,Aa} \\ A_j \to a_i: \quad \Delta n_{i,a} = -1 \quad \mathbb{E}[\Delta n_{i,a}|A_j \to a_i] &= -\frac{1}{Nwj}n_{ij,aA}. \end{aligned} \tag{2}$$

The probability that there is a mutation from the intermediate mutant into the final mutant, which will then cary the mutant lineage to fixation is given by

$$P = \mu\frac{1+s}{Nw}n_a \tag{3}$$

and the expectation is

$$\Delta n_{i,a} = n_{i,A} \quad \mathbb{E}[\Delta n_{i,a}|\text{ second mutation}] = n_{i,A}\mu\frac{1+s}{Nw}n_a, \tag{4}$$

Summing these expectations, we compute the total expected change in the mutant numbers on nodes of fixed degree $i$, $n_{i,a}$, in one update, as

$$\mathbb{E}[\Delta n_{i,a}] = \frac{1}{Nw}\left\{n_{i,A}\mu(1+s)n_a + \sum_j \frac{1}{j}[(1-\mu)(1+s)n_{ij,Aa} - n_{ij,aA}]\right\}. \tag{5}$$

This change in the mutant numbers in one generation, as $N \to \infty$ and $\Delta T = 1/N \to 0$, is

$$\frac{d}{dt} n_{i,a} = \frac{1}{w} \left\{ n_{i,A}\mu(1+s)n_a + \sum_j \frac{1}{j}[(1-\mu)(1+s)n_{ij,Aa} - n_{ij,aA}] \right\}. \tag{6}$$

The dynamics of the mutant on nodes of different degrees therefore critically depend on the frequencies of edge types in the network. To get a full description of the deterministic dynamics, we also need to calculate the change in edge type for one update.

##### 1.1.2 Edge dynamics

In a well-mixed population or on the complete graph, keeping track of the mutant frequencies on all nodes of degree $i$ (in effect, one single frequency) is sufficient for determining the evolutionary fate of the population. However, here, the replacement between mutant and wildtype depends on the frequency of edges that connect the two in network population, in other words how many edges between mutants and the wildtype individuals. For simplicity and clarity, in what follows we assume $s << 1$ and $N\mu << 1$. The edge count $n_{ij,aA}$ can change in two types of events. The first type is a replacement event between a mutant occupying a node of degree $i$ and a wild-type occupying a node of degree $j$. The expected update in $n_{ij,aA}$ is given by

$$\begin{aligned} a_i \to A_j: \quad & \Delta n_{ij,aA} = -1 \quad \mathbb{E}[\Delta n_{ij,aA} | a_i \to A_j] = -\frac{1}{Ni} n_{ij,aA} + o(s) + o(N\mu) \\ A_j \to a_i: \quad & \Delta n_{ij,aA} = -1 \quad \mathbb{E}[\Delta n_{ij,aA} | A_j \to a_i] = -\frac{1}{Nj} n_{ij,aA} + o(s) + o(N\mu). \end{aligned} \tag{7}$$

The combined expected change under these two events is given by

$$\mathbb{E}[\Delta n_{ij,aA} | i \leftrightarrow j] = -\frac{1}{N}\left(\frac{1}{i} + \frac{1}{j}\right) n_{ij,aA} + o(s) + o(N\mu). \tag{8}$$

The second event type is when a third individual occupying a node of degree $k$ replaces a node of degree $i$ or $j$. There are four possible ways for the heterogeneous edge to change in number under this type of event. For example, the probability of $a_k \to A_j a_i$ ( $a_k A_j a_i$ turning into $a_k a_j a_i$), is

$$P(a_k \to A_j a_i) = -\frac{1}{Nk} n_{kj,aA} \frac{n_{kji,aAa}}{n_{kj,aA}(j-1)} + o(s) + o(N\mu). \tag{9}$$

Following similar calculations as above,

$$
\begin{aligned}
a_k \to A_j a_i : \quad \Delta n_{ij,aA} &= -(j-1) \quad \mathbb{E}[\Delta n_{ij,aA} | a_k \to A_j a_i] = -n_{kji,aAa}/(Nk) + o(s) + o(N\mu) \\
A_k \to a_j a_i : \quad \Delta n_{ij,aA} &= +(j-1) \quad \mathbb{E}[\Delta n_{ij,aA} | A_k \to a_j a_i] = +n_{kji,Aaa}/(Nk) + o(s) + o(N\mu).
\end{aligned}
\tag{10}
$$

The expected change is then given by

$$
\mathbb{E}[\Delta n_{ij,aA} | k \to ji] = \sum_k \frac{1}{Nk}(n_{kji,Aaa} - n_{kji,aAa}) + o(s) + o(N\mu). \tag{11}
$$

Similarly,

$$
\begin{aligned}
a_k \to A_i A_j : \quad \Delta n_{ij,aA} &= +(i-1) \quad \mathbb{E}[\Delta n_{ij,aA} | a_k \to A_i A_j] = +n_{kij,aAA}/(Nk) + o(s) + o(N\mu) \\
A_k \to a_i A_j : \quad \Delta n_{ij,aA} &= -(i-1) \quad \mathbb{E}[\Delta n_{ij,aA} | A_k \to a_i A_j] = -n_{kij,AaA}/(Nk) + o(s) + o(N\mu).
\end{aligned}
\tag{12}
$$

The expected change under these two events is given by

$$
\mathbb{E}[\Delta n_{ij,aA} | k \to ij] = \sum_k \frac{1}{Nk}(n_{kij,aAA} - n_{kij,AaA}) + o(s) + o(N\mu). \tag{13}
$$

Summing (8), (11), and (13) allows us to write out the deterministic edge dynamics

$$
\frac{d}{dt} n_{ij,aA} = -\left(\frac{1}{i} + \frac{1}{j}\right) n_{ij,aA} + \sum_k \frac{1}{k}(n_{kji,Aaa} - n_{kji,aAa} + n_{kij,aAA} - n_{kij,AaA}) + o(s) + o(N\mu) \tag{14}
$$

##### 1.1.3 Deterministic mutant equilibrium frequencies

Since we assumed $s << 1$ and $N\mu << 1$, for the neutral dynamics, the quasi-equilibrium point is calculated by setting the right-hand side of equations (6) and (14) to 0 and solving,

$$
\begin{cases}
0 = \sum_j \frac{1}{j}(n_{ij,Aa} - n_{ij,aA}) & \text{node equilibrium,} \\
0 = -\left(\frac{1}{i} + \frac{1}{j}\right) n_{ij,aA} + \sum_k \frac{1}{k}(n_{kji,Aaa} - n_{kji,aAa} + n_{kij,aAA} - n_{kij,AaA}) & \text{edge equilibrium.}
\end{cases}
\tag{15}
$$

We make two key approximations. The first is a pair approximation, termed triple closure (House and Keeling, 2011). This simplifies the dynamics by expressing triple types as edge types and node types

$$n_{ijk,XYZ} = \frac{j-1}{j}\left((1-\phi)\frac{n_{ij,XY}n_{jk,YZ}}{n_{j,Y}} + \phi\frac{n\bar{d}}{ik}\frac{n_{ij,XY}n_{jk,YZ}n_{ik,XZ}}{n_{i,X}n_{j,Y}n_{k,X}}\right) \tag{16}$$

where $\bar{d}$ is the average node degree and $\phi$ is the fraction of triangles in the network. Information on the higher order description of network topology is lost under this assumption, but the number of equations that describes the network dynamics greatly reduce. The second approximation allows us to compute edge dynamics in terms of the node dynamics, thus

$$\begin{cases} n_{ij,AA} = n_{ij}p_{A|i}(1-c_{ij}p_{a|j}), \\ n_{ij,Aa} = n_{ij}c_{ij}p_{A|i}p_{a|j}, \\ n_{ij,aA} = n_{ij}c_{ij}p_{a|i}p_{A|j}, \\ n_{ij,aa} = n_{ij}p_{a|j}(1-c_{ij}p_{A|i}), \end{cases} \tag{17}$$

where $p_{A|i} = \frac{p_{i,A}}{p_i}$ is the probability of type being $A$ conditional on node being of degree $i$. Similarly for $p_{a|i}$. The $c_{ij}$ defines the fraction of $n_{ij}$ that connects the wildtype to the mutant. The $c_{ij}$ are quantities that capture the clustering of mutants due to network structure, leading to a decrease in the number of $Aa$ edges, since they only exist on the boundaries of clusters.

Approximations (16) and (17) allow us to greatly simplify tracking of the equilibrium node frequencies and we can write

$$\begin{aligned} 0 &= \sum_j n_{ij}c_{ij}(p_{a|i}p_{A|j} - p_{A|i}p_{a|j})\frac{1}{j} \\ &= \sum_j n_{ij}c_{ij}[p_{i|a}(1-p_{a|j}) - (1-p_{a|i})p_{a|j}]\frac{1}{j} \\ &= \sum_j n_{ij}c_{ij}(p_{a|i} - p_{a|j})\frac{1}{j}. \end{aligned} \tag{18}$$

This has a solution $p_{a|i} = p_{a|j}$, which means that in the absence of stochastic drift, nodes of different degrees will all have the same mutant frequency. We can use this to define $q_a$

$$q_a = \frac{1}{\mathbb{E}[i^{-1}]}\sum_i p_i p_{a|i}\frac{1}{i}, \quad \text{where} \quad \mathbb{E}[i^{-1}] = \left(\sum_i p_i \frac{1}{i}\right)^{-1}. \tag{19}$$

Notice that, at the equilibrium, $p_{a|i} = p_{a|j}, p_{a|i} = p_{a|j} = q_a$. Also observe that

$$\begin{aligned}\frac{d}{dt}q_a &= \frac{1}{\mathbb{E}[i^{-1}]}\frac{d}{dt}\sum_i p_{i,a}\frac{1}{i} \\ &= \frac{1}{\mathbb{E}[i^{-1}]}\sum_{ij} n_{ij}c_{ij}(p_{a|i}p_{A|j} - p_{A|i}p_{a|j})\frac{1}{ij} \\ &= 0.\end{aligned} \tag{20}$$

This means that $q_a$ is the conserved quantity or the martingale under the neutral dynamics. This shows that any initial mutant frequency on nodes of degree $i$ will reach frequency $q_a$ given enough time.

This expression greatly simplifies the dynamics and shows that we need only track the frequency of mutants occupying nodes of various degrees and the total number of heterogeneous edges in the system. To calculate the edge dynamics, we first simplify equations (8), (11), and (13). Equation (8) becomes

$$\mathbb{E}[\Delta n_{ij,aA}|i \leftrightarrow j] = -\left(\frac{1}{i} + \frac{1}{j}\right) c_{ij}n_{ij}q_a q_A \tag{21}$$

Equation (11) becomes

$$\begin{aligned}&\mathbb{E}[\Delta n_{ij,aA}|k \to ji] \\ &= \sum_k \frac{1}{k}(n_{kji,Aaa} - n_{kji,aAa}) \\ &= \frac{j-1}{j}\sum_k \frac{1}{k}\Bigg((1-\phi)n_{kj,Aa}\frac{n_{ji,aa}}{n_{j,a}} + \phi\frac{n\bar{d}}{ik}\frac{n_{kj,Aa}n_{ji,aa}n_{ik,aA}}{n_{k,A}n_{j,a}n_{i,a}} \\ &\qquad - (1-\phi)n_{kj,aA}\frac{n_{ji,Aa}}{n_{j,A}} - \phi\frac{n\bar{d}}{ik}\frac{n_{kj,aA}n_{ji,Aa}n_{ik,aa}}{n_{k,a}n_{j,A}n_{i,a}}\Bigg) \\ &= \frac{j-1}{j}q_A q_a\sum_k \frac{1}{k}\Bigg((1-\phi)n_{kj}c_{kj}\frac{n_{ji}(1-c_{ji}q_A)}{n_j} + \phi\frac{n\bar{d}}{ik}\frac{n_{kj}c_{kj}n_{ji}(1-c_{ji}q_A)n_{ik}c_{ik}}{n_k n_j n_i} \\ &\qquad - (1-\phi)n_{kj}c_{kj}\frac{n_{ji}c_{ji}q_a}{n_j} - \phi\frac{n\bar{d}}{ik}\frac{n_{kj}c_{kj}n_{ji}c_{ji}n_{ik}(1-c_{ik}q_A)}{n_k n_j n_i}\Bigg) \\ &= \frac{j-1}{j}q_A q_a\sum_k \frac{1}{k}\Bigg((1-\phi)\frac{n_{kj}n_{ji}}{n_j}c_{kj}(1-c_{ji}q_A) + \phi\frac{n\bar{d}}{ik}\frac{n_{kj}n_{ji}n_{ik}}{n_k n_j n_i}c_{kj}(1-c_{ji}q_A)c_{ik} \\ &\qquad - (1-\phi)\frac{n_{kj}n_{ji}}{n_j}c_{kj}c_{ji}p_a - \phi\frac{n\bar{d}}{ik}\frac{n_{kj}n_{ji}n_{ik}}{n_k n_j n_i}c_{kj}c_{ji}(1-c_{ik}q_A)\Bigg) \\ &= \frac{j-1}{j}q_A q_a\sum_k \frac{1}{k}\Bigg((1-\phi)\frac{n_{kj}n_{ji}}{n_j}c_{kj}(1-c_{ji}) + \phi\frac{n\bar{d}}{ik}\frac{n_{kj}n_{ji}n_{ik}}{n_k n_j n_i}c_{kj}(c_{ik}-c_{ji})\Bigg).\end{aligned} \tag{22}$$

Similarly, equation (13) becomes

$$
\begin{aligned}
&\mathbb{E}[\Delta n_{ij,aA}|k \to ij] \\
&\quad = \sum_k \frac{1}{k}(n_{kij,aAA} - n_{kij,AaA}) \\
&\quad = \frac{i-1}{i} \sum_k \frac{1}{k}\Bigg((1-\phi)n_{ki,aA}\frac{n_{ij,AA}}{n_{i,A}} + \phi\frac{n\bar{d}}{jk}\frac{n_{ki,aA}n_{ij,AA}n_{jk,Aa}}{n_{k,a}n_{i,A}n_{j,A}} \\
&\qquad\qquad - (1-\phi)n_{ki,Aa}\frac{n_{ij,aA}}{n_{i,a}} - \phi\frac{n\bar{d}}{jk}\frac{n_{ki,Aa}n_{ij,aA}n_{jk,AA}}{n_{k,A}n_{i,a}n_{i,A}}\Bigg) \\
&\quad = \frac{i-1}{i} q_A q_a \sum_k \frac{1}{k}\Bigg((1-\phi)n_{ki}c_{ki}\frac{n_{ij}(1-c_{ij}q_a)}{n_i} + \phi\frac{n\bar{d}}{jk}\frac{n_{ki}c_{ki}n_{ij}(1-c_{ij}q_a)n_{jk}c_{jk}}{n_k n_i n_j} \\
&\qquad\qquad - (1-\phi)n_{ki}c_{ki}\frac{n_{ij}c_{ij}q_A}{n_i} - \phi\frac{n\bar{d}}{jk}\frac{n_{ki}c_{ki}n_{ij}c_{ij}n_{jk}(1-c_{jk}q_a)}{n_k n_i n_i}\Bigg) \\
&\quad = \frac{i-1}{i} q_A q_a \sum_k \frac{1}{k}\Bigg((1-\phi)\frac{n_{ki}n_{ij}}{n_i}c_{ki}(1-c_{ij}q_a) + \phi\frac{n\bar{d}}{jk}\frac{n_{ki}n_{ij}n_{jk}}{n_k n_i n_j}c_{ki}(1-c_{ij}q_a)c_{jk} \\
&\qquad\qquad - (1-\phi)\frac{n_{ki}n_{ij}}{n_i}c_{ki}c_{ij}q_A - \phi\frac{n\bar{d}}{jk}\frac{n_{ki}n_{ij}n_{jk}}{n_k n_i n_i}c_{ki}c_{ij}(1-c_{jk}q_a)\Bigg) \\
&\quad = \frac{i-1}{i} q_A q_a \sum_k \frac{1}{k}\Bigg((1-\phi)\frac{n_{ki}n_{ij}}{n_i}c_{ki}(1-c_{ij}) + \phi\frac{n\bar{d}}{jk}\frac{n_{ki}n_{ij}n_{jk}}{n_k n_i n_j}c_{ki}(c_{jk}-c_{ij})\Bigg).
\end{aligned}
\tag{23}
$$

Summing up (8), (11), and (13) gives the expected change in edges of node degrees $i$ and $j$ that connect a mutant and a wildtype. Setting this to zero, we write

$$
\begin{aligned}
0 = &-\left(\frac{1}{i}+\frac{1}{j}\right)c_{ij} \\
&+ \frac{j-1}{j}\sum_k \frac{1}{k}\Bigg((1-\phi)\frac{n_{kj}}{n_j}c_{kj}(1-c_{ji}) + \phi\frac{n\bar{d}}{ik}\frac{n_{kj}n_{ik}}{n_k n_j n_i}c_{kj}(c_{ik}-c_{ji})\Bigg) \\
&+ \frac{i-1}{i}\sum_k \frac{1}{k}\Bigg((1-\phi)\frac{n_{ki}}{n_i}c_{ki}(1-c_{ij}) + \phi\frac{n\bar{d}}{jk}\frac{n_{ki}n_{jk}}{n_k n_i n_j}c_{ki}(c_{jk}-c_{ij})\Bigg).
\end{aligned}
\tag{24}
$$

Thus, we find $c_{ij}$ by solving this quadratic system. We then use $c_{ij}$ to express the edge dynamics and find the neutral mutant equilibrium node frequencies.

#### 1.2 Analysis of the stochastic dynamics

In this section, we use the quasi equilibrium results from the previous section to derive the stochastic equations for our model. We show our governing equation is identical to the well-mixed population case with the following changes: an effective selection coefficient $\alpha s$ and an effective extended fixation time $\lambda T_{fix}$. This result leads to approximations for the crossing probability and time, by simply utilizing the transition matrix for the analogous well-mixed system. Lastly, we show that the dynamics of fitness landscape crossing reduce to three simple cases when the population size becomes large.

##### 1.2.1 The diffusion approximation

We analyze the stochastic dynamics by considering how the equilibrium frequency $q_a$ changes when an individual on node of degree $i$ is replaced by the opposite genotype, from a node of degree $j$. The associated changes in $q_a$ are given by

$$\begin{aligned} A_j \to a_i : \quad \Delta q_a &= +\frac{1}{\mathbb{E}[i^{-1}]}\frac{1}{Ni} = +\Delta_i \\ a_j \to A_i : \quad \Delta q_a &= -\frac{1}{\mathbb{E}[i^{-1}]}\frac{1}{Ni} = -\Delta_i . \end{aligned} \tag{25}$$

Therefore, the amount that $q_a$ changes when a mutant on a node of degree $j$ replaces a wildtype on a node of degree $i$, depends on $i$. Therefore, using equation (1), and summing across all possible node degrees $j$, we can write the total probability that $q_a$ increases by $\Delta_i$. We can similarly compute the probability that $q_a$ decreases by $\Delta_i$. The two expressions are

$$\begin{aligned} P\left(\Delta q_{a,i} = +\Delta_i\right) &= (1-\mu)\frac{1+s}{Nw} q_a q_A \sum_j n_{ij} c_{ij} \frac{1}{j} \\ P\left(\Delta q_{a,i} = -\Delta_i\right) &= \frac{1}{Nw} q_a q_A \sum_j n_{ij} c_{ij} \frac{1}{j} . \end{aligned} \tag{26}$$

We also need to compute the frequency change due to mutation. Since we assume that the second mutant fixes immediately, the change in frequency is $q_A$, leading a final frequency of 1. This probability is given by

$$P\left(\Delta q_a = +q_A\right) = \mu \frac{1+s}{w} q_a . \tag{27}$$

Lastly, we have the probability of $q_a$ not changing, which is simply one minus all the probabilities above. This is given by

$$P\left(\Delta q_a = 0\right) = 1 - P\left(\Delta q_a = +q_A\right) - \sum_i P\left(\Delta q_a = \pm\Delta_i\right). \tag{28}$$

Putting it all together, the probability $\Phi(q_a, t)$ of reaching the state of all mutant at time $t$ starting with $q_a$ is described by the backward equation given by (Gardiner, 2009)

$$\Phi(q_a, t + \Delta t) = \sum_{ij} P\left(\Delta q_a = \pm\Delta_i\right)\Phi\left(q_a \pm \Delta_i, t\right) + P\left(\Delta q_a = 0\right)\Phi\left(q_a, t\right) + P\left(\Delta q_a = +q_A\right)\Phi\left(1, t\right) \tag{29}$$

Explicitly writing out the transition probabilities,

$$\begin{aligned}\Phi(q_a,& t + \Delta t) \\ =& \frac{1}{Nw} q_a q_A \sum_{ij} n_{ij} c_{ij} \frac{1}{j}\Bigg[(1-\mu)(1+s)\Phi\left(q_a + \Delta_i, t\right) + \Phi\left(q_a - \Delta_i, t\right)\Bigg] \\ &+ \left(1 - \left[(1-\mu)(1+s)+1\right]\frac{1}{Nw} q_a q_A \sum_{ij} n_{ij} c_{ij} \frac{1}{j} - \frac{\mu(1+s)}{w} q_a\right)\Phi\left(q_a, t\right) + \frac{\mu(1+s)}{w} q_a \Phi\left(1, t\right).\end{aligned} \tag{30}$$

Rearrange the terms, and recall that $\Delta t = 1/N$,

$$\begin{aligned}\frac{\Phi(q_a, t+\Delta t) - \Phi(q_a, t)}{\Delta t}& \\ =& \frac{1}{w} q_a q_A \sum_{ij} n_{ij} c_{ij} \frac{1}{j}\Bigg[\frac{\Phi\left(q_a + \Delta_i, t\right) + \Phi\left(q_a - \Delta_i, t\right) - 2\Phi\left(q_a, t\right)}{\Delta_i^2}\Delta_i^2 \\ &\qquad + (s - \mu - s\mu)\frac{\Phi\left(q_a + \Delta_i, t\right) - \Phi\left(q_a, t\right)}{\Delta_i}\Delta_i\Bigg] \\ &+ \frac{N\mu(1+s)}{w} q_a\left[-\Phi\left(q_a, t\right) + \Phi\left(1, t\right)\right].\end{aligned} \tag{31}$$

Since $\Delta t \to 0$ and $\Delta_i \to 0$ in the limit of large population size, and using the fact that $\Phi(1, t) = 1$, we have

$$\frac{\partial\Phi}{\partial t} = \frac{1}{w} q_a q_A \sum_{ij} n_{ij} c_{ij} \frac{1}{j}\left(\frac{1}{N^2\mathbb{E}[i^{-1}]^2 i^2}\frac{\partial^2\Phi}{\partial p_a^2} + (s - \mu - s\mu)\frac{1}{N\mathbb{E}[i^{-1}]i}\frac{\partial\Phi}{\partial p_a}\right) + \frac{N\mu(1+s)}{w} q_a(1 - \Phi). \tag{32}$$

We rewrite this as

$$\frac{\partial\Phi}{\partial t} = \frac{1}{w} q_a q_A\left(\frac{1}{\lambda N}\frac{\partial^2\Phi}{\partial p_a^2} + \frac{\alpha}{\lambda}(s - \mu - s\mu)\frac{\partial\Phi}{\partial p_a}\right) + \frac{N\mu(1+s)}{w} q_a(1 - \Phi), \tag{33}$$

where

$$\alpha = \mathbb{E}[i^{-1}] \sum_{ij} \frac{c_{ij} n_{ij}}{Nij} \left( \sum_{ij} \frac{c_{ij} n_{ij}}{Ni^2 j} \right)^{-1}, \tag{34}$$

and

$$\lambda = \left( \sum_{ij} \frac{c_{ij} n_{ij}}{Ni^2 j} \right)^{-1} \mathbb{E}[i^{-1}]^2. \tag{35}$$

In the limit of weak selection $s \ll 1$, weak mutation $N\mu \ll 1$, and large population size $1/N \ll 1$,

$$\frac{\partial \Phi}{\partial t} = \frac{1}{\lambda} q_a (1 - q_a) \left( \frac{1}{N} \frac{\partial^2 \Phi}{\partial p_a^2} + \alpha s \frac{\partial \Phi}{\partial p_a} \right) + N\mu q_a (1 - \Phi). \tag{36}$$

Solving for $\Phi(q_a^0)$ gives the probability of fixation at time $t$ when starting with any initial condition $q_a^0$. For a $q_a^0$ corresponding with the mutant appearing in one copy in any node of the graph with equal probability, doing a Taylor expansion, we can write

$$\begin{aligned} \Phi_{q_a^0} &= \sum_i p_i \Phi \left( \frac{1}{\mathbb{E}[i^{-1}]} \frac{1}{iN} \right) \\ &\approx \sum_i p_i \left[ \Phi(0) + \frac{1}{\mathbb{E}[i^{-1}]} \frac{1}{iN} \Phi'(0) \right] \\ &= \sum_i p_i \frac{1}{\mathbb{E}[i^{-1}]} \frac{1}{iN} \Phi'(0) \\ &\approx \Phi \left( \frac{1}{\mathbb{E}[i^{-1}]} \sum_i p_i \frac{1}{iN} \right) \\ &= \Phi \left( \frac{1}{N} \right). \end{aligned} \tag{37}$$

We approximate the crossing time and probability by solving for $\Phi\left(\frac{1}{N}\right)$.

##### 1.2.2 Comparison to the well-mixed model

While the landscape crossing time and probability can be approximated by solving equation (36), here, we can also draw connections to the landscape crossing model in a well-mixed population. This allows us to utilize the rich set of analytical tools and intuitions developed around landscape crossing in well-mixed populations, and helps us understand the crossing behavior in network structured populations.

Consider a well-mixed population with population size $N$ and effective population size $\lambda N$. There is an effective selection $\alpha s$ on the first mutant. The second mutant appears on the background of the first with mutation rate $\mu$ and is assumed to fix upon introduction. There is a distinction between population size and effective population size: effective population size is a proxy for stochastic drift in the system and thus it appears only in the frequency update probability, but not in the mutation probability of the second mutant. The transition matrix for this stochastic system is given by

$$T = \begin{pmatrix} 1 & 0 & & & & 0 \\ P_{1\to 0} & P_{1\to 1} & P_{1\to 2} & & & P_{1\to *} \\ & P_{2\to 1} & \ddots & \ddots & & \vdots \\ & & \ddots & \ddots & & \vdots \\ 0 & 0 & \cdots & \cdots & 1-\mu & \mu \\ 0 & 0 & \cdots & \cdots & 0 & 1 \end{pmatrix} \tag{38}$$

of size $(N+1)\times(N+1)$, where, for $0 < i < N$,

$$\begin{cases} P_{i\to i-1} = \frac{1}{\lambda}\frac{1}{w'}\frac{i}{N}\frac{N-i}{N}, \\ P_{i\to i} = 1 - \frac{1}{\lambda}\frac{2+\alpha s}{w'}\frac{i}{N}\frac{N-i}{N} - \mu\frac{1+\alpha s}{w}\frac{i^2}{N^2}, \\ P_{i\to i+1} = (1-\mu)\frac{1}{\lambda}\frac{1+\alpha s}{w'}\frac{i}{N}\frac{N-i}{N}, \\ P_{i\to *} = \mu\frac{1+\alpha s}{w'}\frac{i}{N}. \end{cases} \tag{39}$$

Here, $w'$ is the mean fitness $(N + i\alpha s)/N$. The first $N$ states correspond to the number of the first mutants in the population. State * corresponds to the appearance of the second mutant. The probability $\Phi(i, T)$ of reaching the state of mutant fixation at time $T$, starting with $i$ in the population, is described by the backward equation given by

$$\Phi(i, T+1) = P_{i\to i+1}\Phi\left(i+1, T\right) + P_{i\to i}\Phi\left(i, T\right) + P_{i\to i-1}\Phi\left(i-1, T\right) + P_{i\to *}\Phi(*, T). \tag{40}$$

Substituting $x = i/N, \Delta x = 1/N, t = T/N$, and $\Delta t = 1/N$

$$\Phi(x, t+\Delta t) = P_{x\to x+\Delta x}\Phi\left(x+\Delta x, t\right) + P_{x\to x}\Phi\left(x, t\right) + P_{x\to x-\Delta x}\Phi\left(x-\Delta x, t\right) + P_{x\to *}\Phi(*, t). \tag{41}$$

Explicitly writing out the transition probabilities

$$\begin{aligned}\Phi(x,t+\Delta t)& \\ =&\frac{1}{\lambda N w'}x(1-x)\Big[(1-\mu)(1+\alpha s)\Phi\left(x+\Delta x,t\right)+\Phi\left(x-\Delta x,t\right)\Big] \\ &+\left(1-(1+(1-\mu)(1+\alpha s))\frac{1}{\lambda N w'}x(1-x)-\frac{\mu(1+\alpha s)}{w}x\right)\Phi\left(x,t\right)+\frac{\mu(1+\alpha s)}{w'}x\Phi\left(*,t\right).\end{aligned} \tag{42}$$

Rearranging the terms,

$$\begin{aligned}\frac{\Phi(x,t+\Delta t)-\Phi(x,t)}{\Delta t}& \\ =&\frac{1}{\lambda w'}x(1-x)\left[\frac{\Phi\left(x+\Delta x,t\right)+\Phi\left(x-\Delta x,t\right)-2\Phi\left(x,t\right)}{(\Delta x)^2}(\Delta x)^2\right. \\ &\left.+\left(\alpha s-\mu+\alpha s\mu\right)\frac{\Phi\left(x+\Delta x,t\right)-\Phi\left(x,t\right)}{\Delta x}\Delta x\right] \\ &+\frac{\mu(1+\alpha s)}{w'}x\left[-\Phi\left(x,t\right)+\Phi\left(*,t\right)\right].\end{aligned} \tag{43}$$

Since $\Delta t\to 0$ and $\Delta x\to 0$ in the limit of large population size, and using the fact that $\Phi(*,t)=1$, we have

$$\frac{\partial\Phi}{\partial t}=\frac{1}{\lambda w'}x(1-x)\left[\frac{1}{N}\frac{\partial^2\Phi}{\partial x^2}+(\alpha s-\mu+\alpha s\mu)\frac{\partial\Phi}{\partial x}\right]+\frac{N\mu(1+\alpha s)}{w'}x\left(1-\Phi\right). \tag{44}$$

In the limit of weak selection $s\ll 1$, weak mutation $N\mu\ll 1$, and large population size $1/N\ll 1$,

$$\frac{\partial\Phi}{\partial t}=\frac{1}{\lambda}x(1-x)\left(\frac{1}{N}\frac{\partial^2\Phi}{\partial x^2}+\alpha s\frac{\partial\Phi}{\partial x}\right)+N\mu x\left(1-\Phi\right). \tag{45}$$

This shows that in the limit of weak selection, the crossing dynamics for a network population mimics the crossing dynamics of a well-mixed system with modified selection and drift. In this context, the arsenal of tools developed for well-mixed evolutionary theory can be applied to study landscape crossing in a network population. One such application is finding the crossing time and probability using Markov chain theory. Since equation (45) is identical to (36), and (45) is the continuous approximate of Markov system described by (38), we can use this Markov system as a proxy for landscape crossing dynamics on a network. As an alternative to solving (36) numerically, this proxy allows us efficiently calculate the crossing probability by finding the left eigenvector of $T$ with an eigenvalue of 1 (Karlin, 2014; Nowak, 2006). The crossing probability, assuming the number of initial intermediate mutants is one, corresponds to the second entry of the eigenvector. Similarly, fixation time is given by taking the first row of $(I - T[1:N, 1:N])^{-1}$, dot product it with the left eigenvalue of $T$, and divide by the probability of fixation (Hindersin et al., 2016).

##### 1.2.3 Continuous time branching point process approximation

As $N$ becomes large the tunneling probability approaches the approximation from continuous time branching process (Weissman et al., 2009). This probability is given by

$$\Phi = \frac{\alpha s - \lambda\mu + \sqrt{(\alpha s + \lambda\mu)^2 + 4\lambda\mu}}{2(1 + \alpha s)}. \tag{46}$$

Under weak selection $|Ns| << 1$, eq. (46) becomes

$$\Phi \approx \frac{\alpha s - \lambda\mu + \sqrt{(\alpha s + \lambda\mu)^2 + 4\lambda\mu}}{2}. \tag{47}$$

If selection dominates the mutation rate , $|s| \gg 2\sqrt{\mu}$, we Taylor expand eq. 47 with respect to $\mu$. This leads to

$$\Phi = \frac{1}{2}\left[\alpha s + \alpha|s| - \lambda\mu + \frac{2\lambda}{\alpha|s|}\mu + o(s^2)\right]. \tag{48}$$

For a beneficial intermediate mutation, the leading term is

$$\Phi \approx \alpha s. \tag{49}$$

For a deleterious intermediate mutation, the leading term is

$$\begin{aligned}\Phi &\approx -\frac{1}{2}\lambda\mu + \frac{\lambda\mu}{\alpha|s|}\\ &\approx \frac{\lambda\mu}{\alpha|s|}.\end{aligned} \tag{50}$$

If the mutation rate dominates selection, $|s| \ll 2\sqrt{\mu}$, we Taylor expand eq. 47 with respect to $s$. This leads to

$$\begin{aligned}\Phi &= \frac{1}{2}\left[-\lambda\mu + \sqrt{(\lambda\mu)^2 + 4\lambda\mu} + o(s)\right]\\ &\approx \sqrt{\lambda\mu},\end{aligned} \tag{51}$$

by taking the leading term in the Puiseux expansion in $\mu$. The above limiting behaviors are summarized as

$$\Phi = \begin{cases} \frac{\lambda\mu}{\alpha|s|} & \text{for } s \ll -2\sqrt{\mu} \\ \sqrt{\lambda\mu} & \text{for } -2\sqrt{\mu} \ll s \ll 2\sqrt{\mu} \\ \alpha s & \text{for } s \gg 2\sqrt{\mu}. \end{cases} \tag{52}$$

Since this approximation neglects tunneling through fixation of the first mutation if it is neutral, we add a small correction term of $1/N$ that leads to

$$\Phi = \begin{cases} \frac{\lambda\mu}{\alpha|s|} & \text{for } s \ll -2\sqrt{\mu} \\ \sqrt{\lambda\mu} + \frac{1}{N} & \text{for } -2\sqrt{\mu} \ll s \ll 2\sqrt{\mu} \\ \alpha s & \text{for } s \gg 2\sqrt{\mu}. \end{cases} \tag{53}$$

**Figure S1A** shows that the approximation in equation (53) provides accurate prediction of the ratio of crossing probability calculated directly from the state transition matrix in the deleterious, neutral, and beneficial regime. For smaller population sizes, the deleterious intermediate can be carried to fixation through drift. A population structure with an amplification factor lower than one decreases the fixation probability, leading to a higher crossing rate. This introduces non-monotonicity in the ratio of crossing probability that vanishes with increasing $N$ (**Figure S1B**). As a result, when $N$ increases, the equation ratio of crossing probability calculated directly from the state transition matrix converges to equation (52). Increasing the mutation rate also reduce the non-monotonicity in the ratio of crossing probability (**Figure S1C**). This is due to the fact that the tunneling probability is an increasing function of the mutation rate, making stepwise fixation a less significant contributor to the crossing probability.

### 2 Landscape crossing results on various graphs families

Our results generalize to all network families of increasing complexity. Here we test four additional network families. The first graph family we consider is the random $k$-regular graph. $k$-regular graphs are isothermal since every node has $k$-edges. The amplification factor is therefore 1. In contrast, the degree $k$ and the fraction of triangles $\theta$ influence the fixation time for the case of a single mutation entering the population. $k$-regular graphs are an ideal network family for studying the time of fixation on landscape crossing. Here we use degree preserving edge swap to tune the number of triangles in the graph, as outlined in (Kuo and

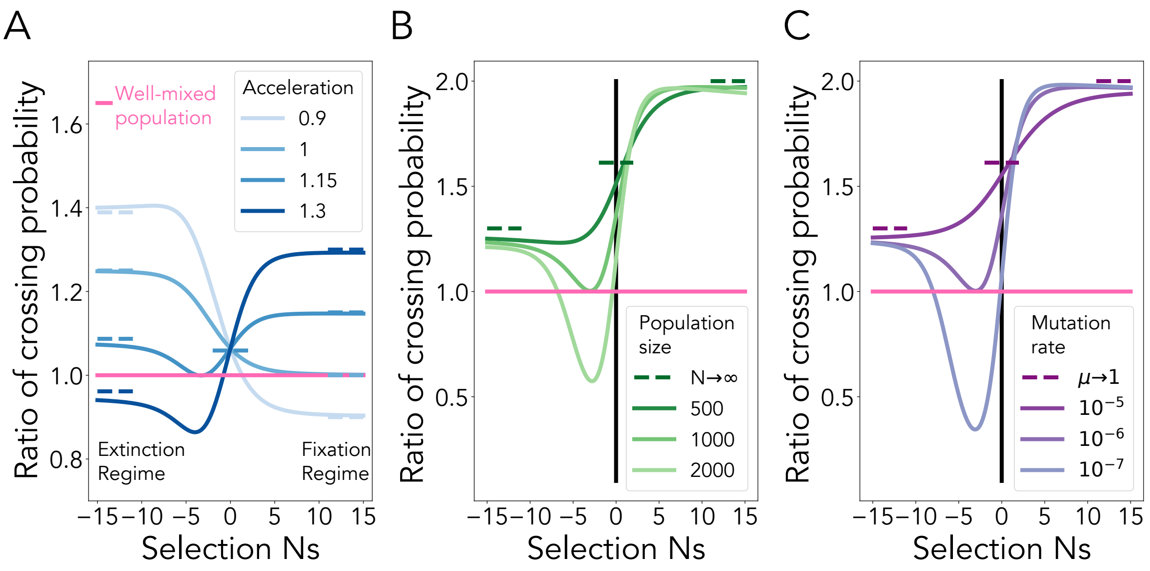

Figure S1: **The continuous time branching process accurately describes the dynamics of crossing in the large population limit.** Here, the solid line is the ratio of tunneling rate between a structured population and the well-mixed population calculated from the state transition matrix in 38. **A,** The dashed lines represent the limiting behavior from equation 53. The population size is 1000. The acceleration factor is fixed at 1.25. The mutation rate is $10^{-6}$. **B,** The dashed lines represent the limiting behavior from equation 52. The amplification factor is 2 and acceleration factor is 2.6. The mutation rate is $10^{-6}$. **C,** The dashed lines represent the limiting behavior from equation 52. The amplification factor is 2 and acceleration factor is 2.6. The population size is 1000.

Carja, 2021). The acceleration factor is given by

$$\lambda = \left(1 - \frac{1}{(k-1)(1-\theta)}\right)^{-1}. \tag{54}$$

Decreasing the mean degree and increasing the fraction of triangles leads to an increase in the crossing probability of fitness valleys and plateaus (**Figure S2**). Intuitively, decreasing the mean degree and increasing the fraction of triangles causes the time to fixation/extinction to also increase. This provides a longer time for the intermediate mutation to persist in the population and acquire the second mutation to carry the mutant lineage to fixation.

The second graph family we consider is the small-world family of networks, which exhibit the small-world effect, observed in many social networks (Watts and Strogatz, 1998). The network generation algorithms for these graphs depend on two parameters: the first parameter controls the degree of the cyclic graph the network is initialized to and the second parameter controls the rewiring probability of the cyclic graph. We calculate the amplification and acceleration factors for these networks using $10^7$ simulations of single mutant

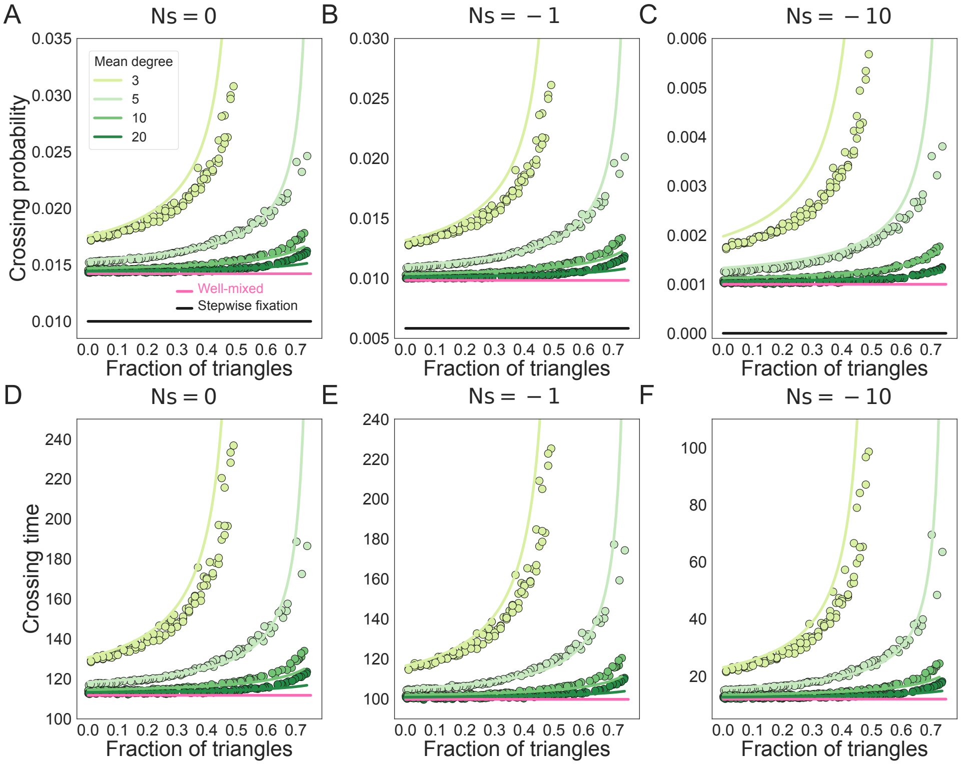

Figure S2: **Lower mean degree and higher triangle count increase the crossing probability and crossing time of fitness plateaus and valleys.** The dots represent the probability and time of acquiring a second mutation calculated from ensemble averages across $10^7$ replicate Monte Carlo simulations. The black line represents the probability of acquiring a second mutation assuming sequential fixation. The solid colored lines represent the fitness valley crossing probabilities and times for the network-structured population, calculated from the state transition matrix in (38). Here the degree distribution is held constant as we vary the fraction of triangles in the graphs, N=100, and mutation rate is $10^{-4}$. The color indicates the mean degree of the network as in the legend. The fraction of triangles in the graph is tuned using edge swapping operations. Panels **A** and **D**, $s = 0$. Panels **B** and **E**, $s = -0.01$. Panels **C** and **F**, $s = -0.1$.

fixation, and we fit these two evolutionary descriptors for the network to network generation parameters, using cubic splines. The tunneling probability and time is calculated from the fitted function and plotted on top of simulations results in (**Figure S3**). The crossing probability and time decreases with increasing mean degree, similar to $k$-regular graphs. As the rewiring probability increases, the graphs begin to exhibit the "small-world" effect where the diameter of the graph decreases and the nodes in the graph are more accessible to one another. Mutants spread faster in the network, decreasing the fixation/extinction time and leading to a lower crossing probability.

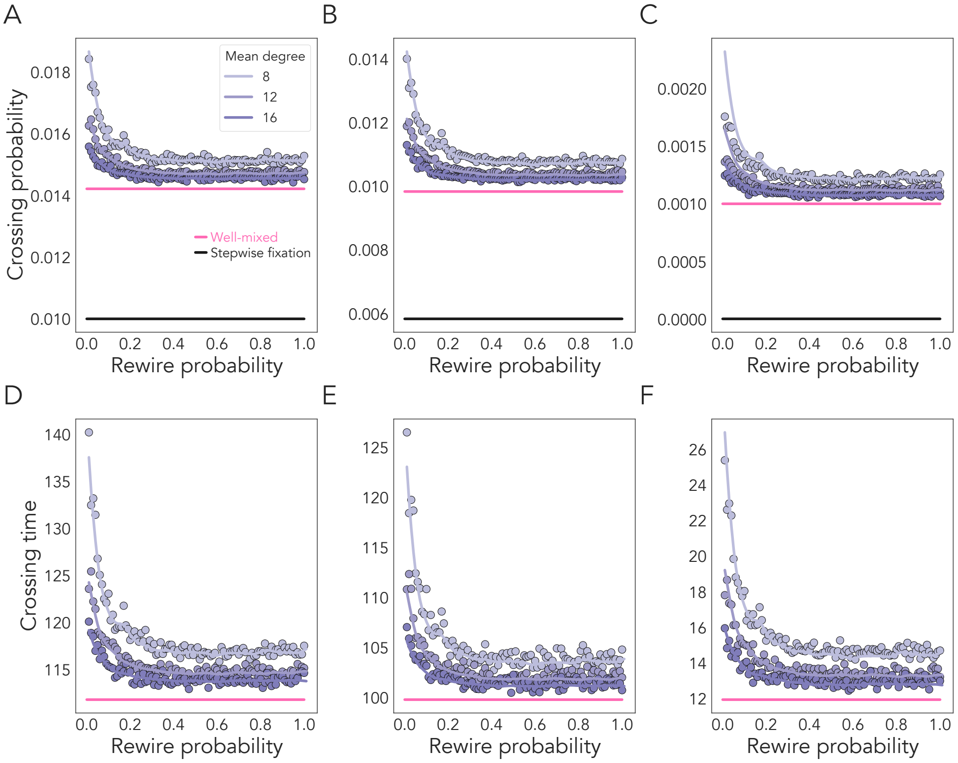

Figure S3: **Probabilities and times for fitness valley crossing for small world graphs.** Here, the graph size is $N = 100$ and mutation rate $\mu = 10^{-4}$. The dots represent ensemble averages across $10^7$ replicate Monte Carlo simulations. The black line represents the probability of acquiring a second mutation assuming sequential fixation. The solid colored lines represent the crossing probability (top) and time (bottom) for small world network-structured populations, using the state transition matrix in (38). Panels **A** and **D**, $s = 0$. Panels **B** and **E**, $s = -0.01$. Panels **C** and **F**, $s = -0.1$.

We next consider random geometric graphs (Waxman, 1988; Penrose et al., 2003). In this family of graphs, nodes have spatial positions randomly drawn from a probability distribution to model spatially homogeneous populations (using the uniform distribution) or populations with heterogeneous spatial density (using the normal distribution). Once the spatial locations of the nodes are determined, the generating algorithm iterates through all pairs of nodes. An edge is created between two nodes if the pair-wise distance is below some prescribed threshold. Here, we show that the network topology increases crossing probability for plateaus and valleys when the network connectivity is low (**Figure S4**).

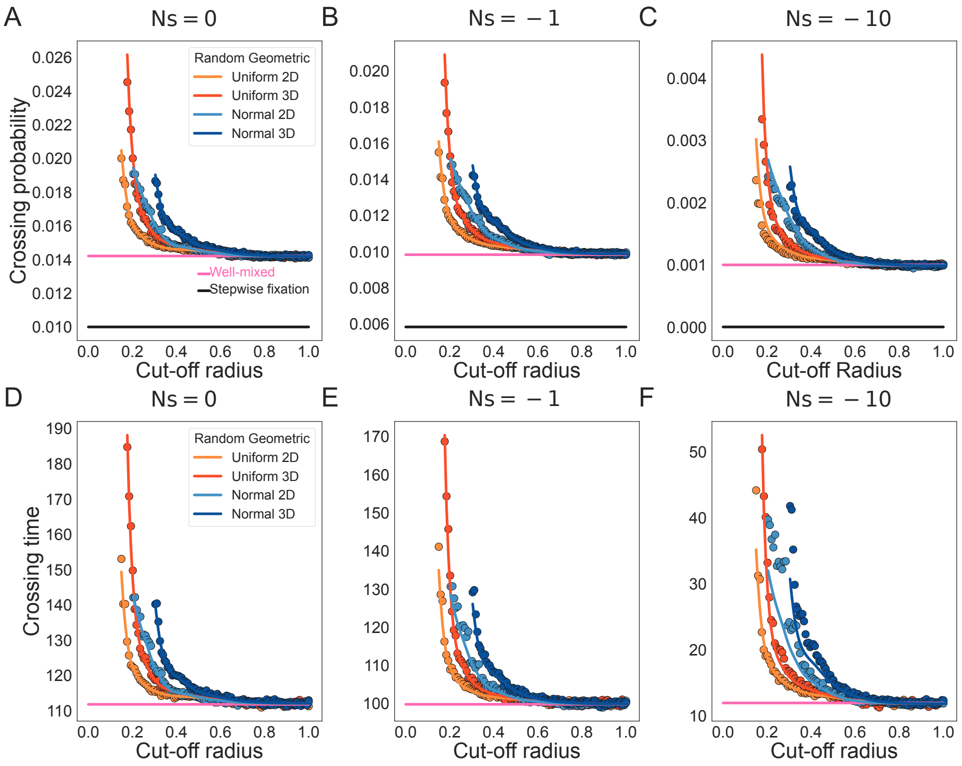

Figure S4: **Landscape crossing probabilities and times for random geometric graphs.** Here, the graph size is $N = 100$ and the mutation rate $\mu = 10^{-4}$. The dots represent ensemble averages across $5 \times 10^6$ replicate Monte Carlo simulations. The colors of the dots represent geometric graphs generated using different probability distributions. The random points of the graphs are kept constant within the same color graph, but are different in the cut-off radius that determines the connectivity of the network, plotted on the x-axis. The black line represents the probability of acquiring a second mutation assuming sequential fixation. The solid colored lines represent the crossing probability (top) and time (bottom) for geometric network-structured populations, using the state transition matrix in (38). Panels **A** and **D**, $s = 0$. Panels **B** and **E**, $s = -0.01$. Panels **C** and **F**, $s = -0.1$.

We next directly study the role of the graph network amplification factor on the crossing dynamics. In **Figure S5**, we use random geometric graphs with uniform and normal distribution of spatial points, small world graphs and Erdoes Renyi graphs, and tune the amplification factor through changing the mixing pattern of these graphs using methods from Kuo et al. (2021). The method changes the mixing pattern while keeping the degree distribution constant. Since degree distribution greatly affects fixation time, and acceleration factor by extension, this allows us to tune the amplification parameter with minimal disturbance to the acceleration factor (Kuo and Carja, 2021) However, this method does not completely decouple amplification factor and acceleration factor. **Figure S5** shows the counter-intuitive result that increasing the amplification factor of a network leads to an increased crossing rate for neutral and deleterious intermediate mutations. This is because the ratio of acceleration and amplification factor is the determinant in crossing probability under valley crossing. The increase in amplification is accompanied by an increase in acceleration. If the increase in the acceleration factor is greater than the increase in the amplification factor, the fitness valley crossing probability would end up increasing. Further investigation into tunable network parameters that allows independent adjustment of the amplification factor is needed to numerically investigate its effect.

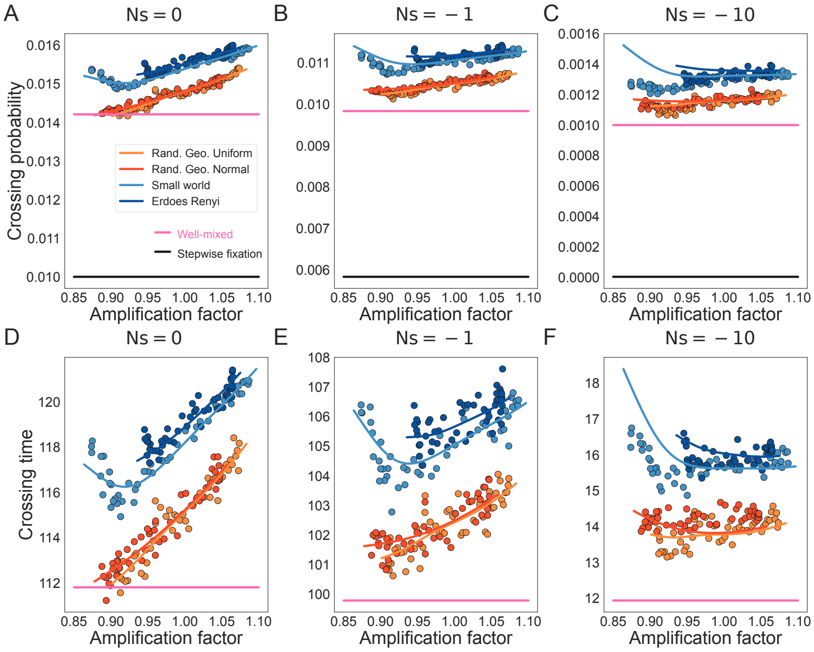

Figure S5: **Exploring the effects of amplification parameters in landscape crossing.** Here, graph size is $N = 100$ and mutation rate is $\mu = 10^{-4}$. The dots represent ensemble averages across $5 \times 10^6$ replicate Monte Carlo simulations. The colors of the dots represent graphs from different graph families as in the legend. The amplification parameter is altered through changing network connectivity pattern. The degree distribution is kept constant for graphs plotted in the same color. The solid colored lines represent the crossing probability (top) and time (bottom) for geometric network-structured populations, using the state transition matrix in (38). Panels **A** and **D**, $s = 0$. Panels **B** and **E**, $s = -0.01$. Panels **C** and **F**, $s = -0.1$.

### 3 Application to the bone marrow stem cell population architectures. Rates of neoplasm initiation.

We apply our model to study the propagation of somatic mutations in the stem cell architectures of the bone marrow to understand spatial factors shaping rates of leukemia initiation. We use networks inferred from quantitative imaging of hematopoietic stem cell niche in the bone marrow Kuo et al. (2021); Coutu et al. (2018); Gomariz et al. (2018). The number of nodes from each image ranges from 300-3000 cells. The amplification and acceleration factors of the bone marrow network structure is calculated from the Monte Carlo simulation of single mutation fixation of $10^7$ replicate simulations at $Ns = 0.01$. The two factors are used to derive the analytical approximation for the crossing probability in these cellular populations, shown here together with simulations (**Figure S6A**). Although the image data provides precise spatial description of the haematopoietic stem cell niches, these only represent a small portion of the HSC population. Previous models estimate the number of haematopoietic stem cells that are actively making white blood cells at any one time to be in the range of 50,000-200,000 (Lee-Six et al., 2018). Here, we assume the overall spatial distribution of the HSC population across the bone marrow is identical to our sample. We use linear regression to extrapolate the amplification factor to the lower bound population of 50,000 (**Figure S6B**). The acceleration factors inferred from data are close to 1, so we assume the larger population also has acceleration factor of 1.

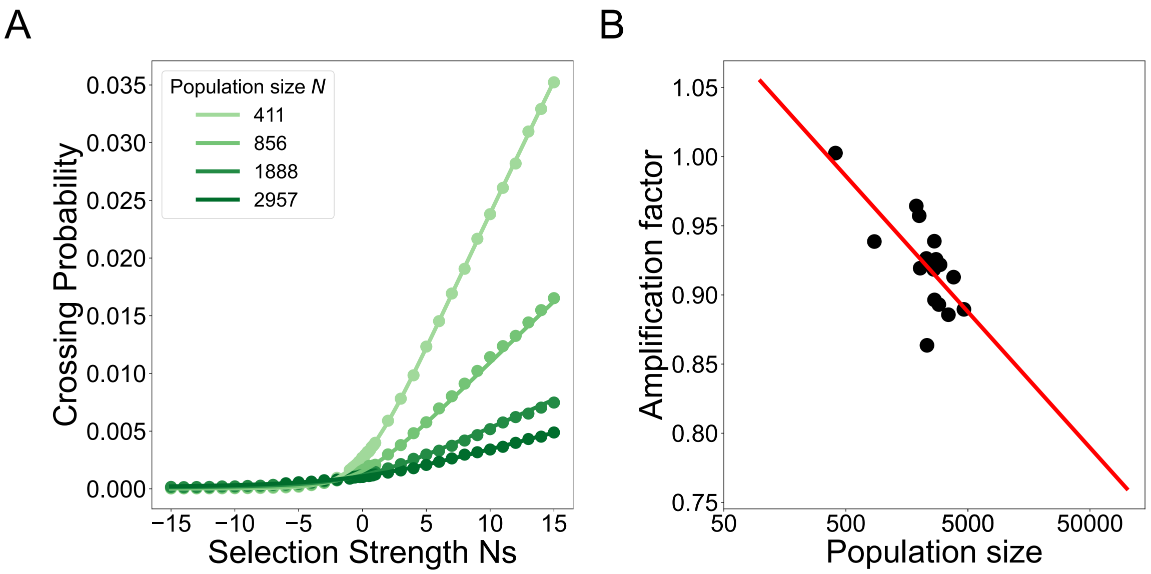

Figure S6: **A two hit model for leukemia development in the bone marrow. Panel A** shows the probability of acquiring two mutations, assumed to lead to initiation of leukemia. Here, the dots represent probability calculated from $10^7$ simulations using population structures inferred from 4 different bone marrow fluorescence microscopy images. The colors represents different bone marrow samples and the number of niche components detected in each sample. Mutation rate is assumed to be $\mu = 10^{-6}$. **Panel B** shows the amplification parameter calculated from $10^7$ simulations of single mutation fixation. The dots represent population structures inferred from different bone marrow samples. The line represents the linear regression log population size and amplification factor. Increased population size correlates with increased suppression of selection.

Crispin Gardiner. *Stochastic methods*, volume 4. Springer Berlin, 2009.

Alvaro Gomariz, Patrick M Helbling, Stephan Isringhausen, Ute Suessbier, Anton Becker, Andreas Boss, Takashi Nagasawa, Grégory Paul, Orcun Goksel, Gábor Székely, et al. Quantitative spatial analysis of haematopoiesis-regulating stromal cells in the bone marrow microenvironment by 3d microscopy. *Nature Communications*, 9(1):1–15, 2018.

Laura Hindersin, Marius Möller, Arne Traulsen, and Benedikt Bauer. Exact numerical calculation of fixation probability and time on graphs. *Biosystems*, 150:87–91, 2016.

Thomas House and Matt J Keeling. Insights from unifying modern approximations to infections on networks. *Journal of The Royal Society Interface*, 8(54):67–73, 2011.

Samuel Karlin. *A first course in stochastic processes.* Academic press, 2014.

Yang Ping Kuo and Oana Carja. Evolutionary graph theory beyond pairwise interactions: higher-order network motifs shape times to fixation in structured populations. *bioRxiv*, 2021. doi: 10.1101/2021.06.26.450017. URL `https://www.biorxiv.org/content/early/2021/06/28/2021.06.26.450017`.

Yang Ping Kuo, César Nombela Arrieta, and Oana Carja. A theory of evolutionary dynamics on any complex spatial structure. *bioRxiv*, 2021. doi: 10.1101/2021.02.07.430151. URL `https://www.biorxiv.org/content/early/2021/02/08/2021.02.07.430151`.

Henry Lee-Six, Nina Friesgaard Øbro, Mairi S Shepherd, Sebastian Grossmann, Kevin Dawson, Miriam Belmonte, Robert J Osborne, Brian JP Huntly, Inigo Martincorena, Elizabeth Anderson, et al. Population dynamics of normal human blood inferred from somatic mutations. *Nature*, 561(7724):473–478, 2018.

Martin A Nowak. *Evolutionary dynamics: exploring the equations of life.* Harvard university press, 2006.

Mathew Penrose et al. *Random geometric graphs*, volume 5. Oxford university press, 2003.

Duncan J Watts and Steven H Strogatz. Collective dynamics of small world networks. *Nature*, 393(6684): 440–442, 1998.

Bernard M Waxman. Routing of multipoint connections. *IEEE journal on selected areas in communications*, 6(9):1617–1622, 1988.

Daniel B Weissman, Michael M Desai, Daniel S Fisher, and Marcus W Feldman. The rate at which asexual populations cross fitness valleys. *Theoretical Population Biology*, 75(4):286–300, 2009.
